# Value-based decisions involve sequential sampling from memory

**DOI:** 10.1101/269290

**Authors:** Akram Bakkour, Ariel Zylberberg, Michael N. Shadlen, Daphna Shohamy

## Abstract

Deciding between two equally appealing options can take considerable time. This observation has puzzled economists and philosophers, because more deliberation only delays the reward. Here we show that this seemingly irrational behavior is explained by the constructive use of memory. Using functional brain imaging in humans, we show that how long it takes to decide between two familiar food items is related to activity in the hippocampus, within specific regions shown to be associated with the retrieval of long-term memories. Moreover, we show that value is partially constructed during deliberation to resolve preference, and this constructive process changes behavior and brain responses. These results render memory as a supplier of evidence in value-based decisions, resolving a central paradox of choice.

## Main Text

Some decisions take more time than others. A common but puzzling phenomenon is that even seemingly simple decisions between familiar items whose value is known take more time when they involve a choice between options of similar value. One explanation for why such decisions take more time is that a commitment to a choice depends on the accumulation of evidence to a threshold, and when the evidence is weaker, more samples are required to achieve this criterion. This simple observation holds across many kinds of decisions, whether they are based upon perception of the environment — is the banana green or yellow^1–4^? — or from internal values and preferences — do I prefer a fresh banana or a ripe one^5–8^? For decisions based on perception, it is clear why more time might provide more samples of evidence and therefore improve the accuracy of the decision^9–14^, but for decisions about internal values and preferences, it is unclear what is the source of the evidence and why more samples should be required to decide between options that are close in value.

We investigated where the time goes in value-based decisions, asking where in the brain information processing corresponds to decision time, how this differs from decisions based on perception, and what are the implications for how value is constructed. Our central hypothesis is that value-based decisions exploit hippocampal-dependent memory mechanisms to derive internal evidence bearing on choice preference. Items of similar value require more evidence to resolve the preference, and therefore take more time. Notably, we propose that this process contributes even to decisions between items that do not appear at first glance to depend on memory, such as decisions between two highly familiar food items^15^. Further, we hypothesize that this deliberative process also makes the values of the items susceptible to adjustment. Any internal source of evidence bearing on a choice between two items must derive from some stored association between object and value^16–18^. We focused on hippocampal memory systems, because the act of comparing items seems to invite consideration of features that are not captured by a simple object-value association. Such a process is likely to involve memory retrieval, but also episodic simulation and prospection — all of which have been shown to depend on the hippocampus^19–25^. It is this memory-based constructive process that we hypothesized is positioned to supply evidence during deliberation.

To test this hypothesis, we designed a task in which human participants were asked to make a series of choices between two familiar food items (*value-based decisions*) while being scanned with fMRI (**Figure 1**). The subjective value of each individual item was determined for each participant in advance (see *Methods*), so that we could systematically vary the difference in value between the two items (i.e. ΔValue) during the decision task (see also^5, 6^). This allowed us to examine the timing of value-based decisions, its relation to ΔValue, and the brain regions for which activity co-varied with decision time. Because there are many different reasons why brain activity might covary with decision time, the same participants also participated in a control condition that required them to make *perceptual decisions* about the dominant color of random dots on the screen. Finally, to link decision-related activity directly to memory, participants also performed a separate *memory recognition task* that served as a functional localizer to independently identify regions of the brain involved in memory retrieval and to probe their role in value-based versus perceptual decisions.

**Figure 1:**
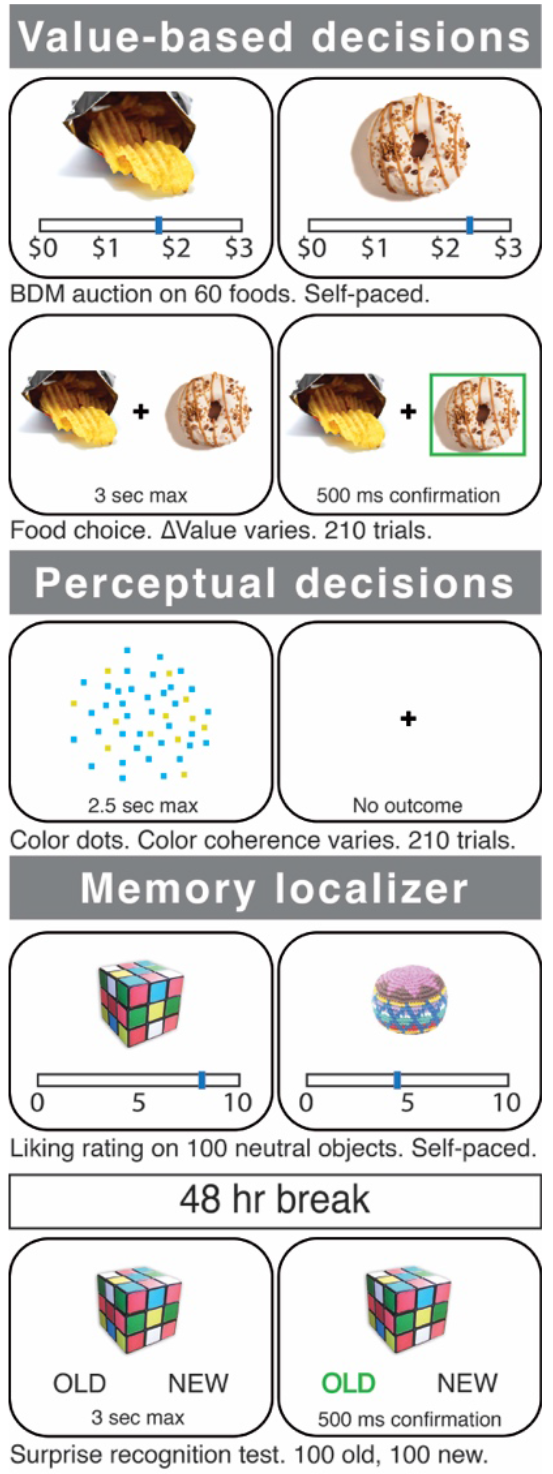
Experimental procedure for evaluating where the time goes in value-based versus perceptual decisions and decision time’s relation to memory. Healthy participants were scanned with fMRI during three different tasks: a value-based decision task (top), a perceptual decision task (middle), and a memory recognition task (bottom). In the value-based decision task, participants were presented with 210 pairs of foods that differed on ΔValue (based on a pre-task auction procedure for rating the items; see *Methods*). Participants were told to choose the item that they preferred and that their choice on a randomly selected trial would be honored at the end of the experiment. In the perceptual decision task, participants were presented with 210 trials of a cloud of flickering blue and yellow dots that varied in the proportion of blue versus yellow (color coherence). Participants were told to determine whether the display was more blue or more yellow. In the recognition memory task, participants underwent a standard memory recognition task using incidental encoding of everyday objects: first, they rated 100 objects (outside of the scanner) and 48 hours later were presented with a surprise memory test in the scanner. The memory test presented participants with the same ‘old’ objects intermixed with 100 ‘new’ objects, one at a time, and asked participants to report their memory for each by indicating whether the presented object was “old” or “new”.

In the value-based and perceptual decision tasks, we expected that decision time would be longer when the choice options were closer in value or when the proportion of blue and yellow dots was closer to 1/2, respectively, but that these decisions would be resolved using different brain mechanisms. Specifically, we predicted that hippocampal BOLD activity would correlate with reaction time (RT) during value-based decisions more so than during perceptual decisions and that this effect would overlap with regions of the hippocampus that show activity related to memory retrieval, independently identified. We also reasoned that memory retrieval during deliberation is a constructive process that may lead to updating of value with consequences for subsequent decisions, as detailed below.

We found that participants made more accurate perceptual decisions when the color was more strongly biased toward blue or yellow (**Figure 2a**, top), and they made decisions more consistent with their subjective valuation when ΔValue was larger (**Figure 2b**, top). In both tasks, decisions between options that are more difficult to discriminate (i.e. ΔValue or color coherence near zero) led to more variable choices (i.e. near chance) and also took longer to complete (**Figure 2**, bottom). We found that choices and reaction time were well-described by a drift diffusion model, which is commonly used to study this sort of decision process (see dark lines in **Figure 2;** for model parameter estimates, see **Supplementary Table 1**, and for individual participant fits see **Supplementary Figure 1**). This is observed in many perceptual and cognitive decisions^26–28^ and is suggestive of a decision process involving sequential sampling and optional stopping based on the level of evidence.

**Figure 2:**
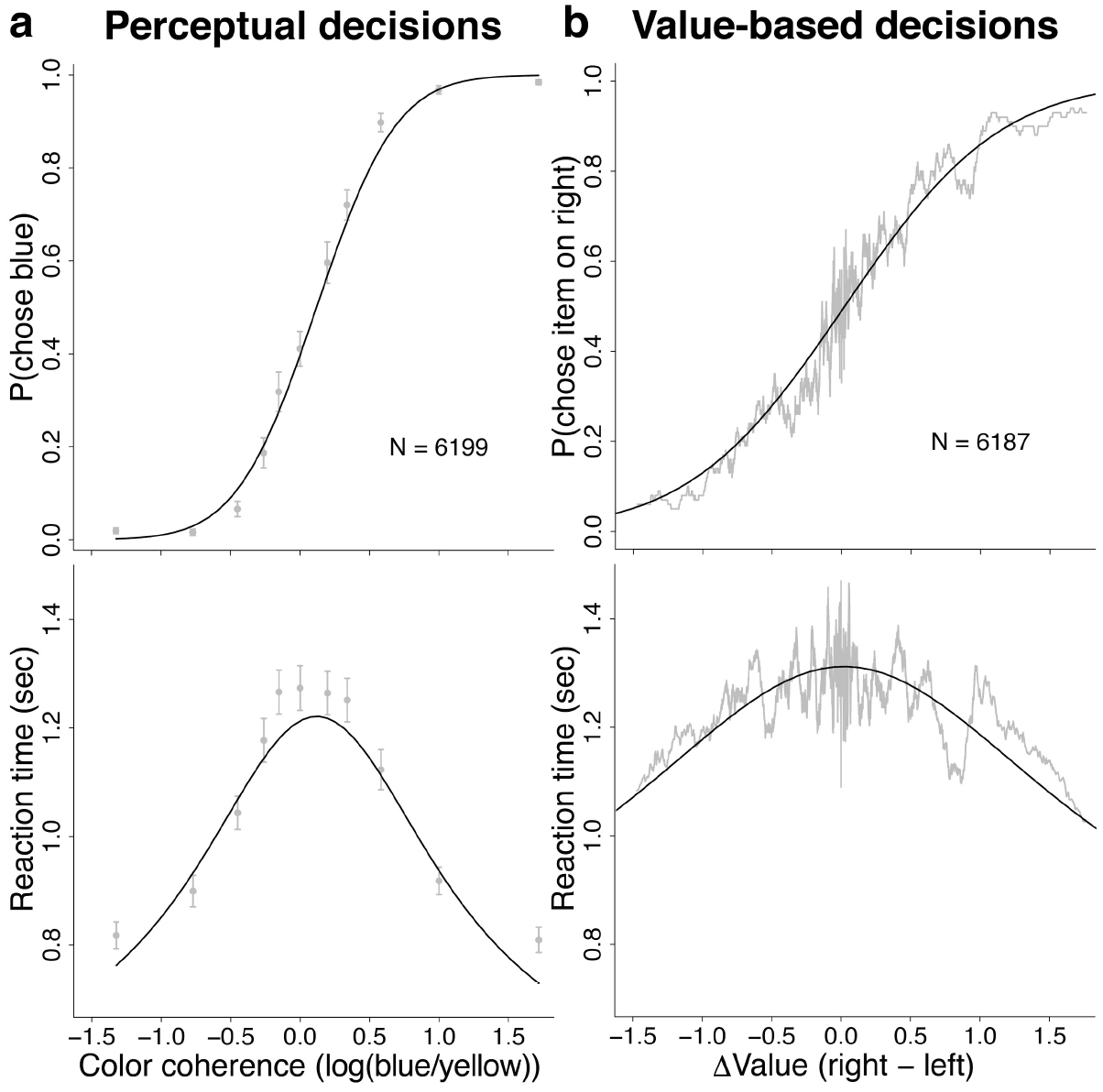
Choices between options that are similar take more time for both perceptual and value-based decisions. Behavioral results for choices (top) and RTs (bottom) for (**a**) perceptual and (**b**) value-based decisions. **a)** The x-axis is the log odds that a dot in the stimulus is blue (color coherence, positive values indicate more blue and negative values indicate more yellow). **b)** The x-axis is the difference in value (from the auction) for the item on the right minus the value of the item on the left side of the screen (ΔValue). Gray symbols are means (error bars are s.e.m.). Gray lines are the running means. Solid black lines are the model fits.

For the color task, the sequence of evidence samples was supplied by the dynamic stimulus, whereas for the value task, the stimuli (and external sources of evidence) were static, leaving open the question of what constituted the evidence being sampled. To determine whether memory processes supply the sequence of samples that support choice and RT during value-based decisions, we conducted a whole-brain analysis to identify regions in the brain that show both (i) a memory retrieval effect (using the separate object-memory localizer task, see *Methods*, **Supplementary Figure 2** and **Supplementary Table 2**) and (ii) an effect of reaction time on BOLD for the value-based task more than for the perceptual task. As shown in **Figure 3a** (and **Supplementary Table 3**), we found a conjunction of these two effects in the hippocampus. This finding shows that BOLD activity in memory-related hippocampal regions was more positively correlated with RT in the value task than in the perceptual task, suggesting a possible role for memory in accounting for the timing of value-based decisions but not perceptual decisions.

**Figure 3:**
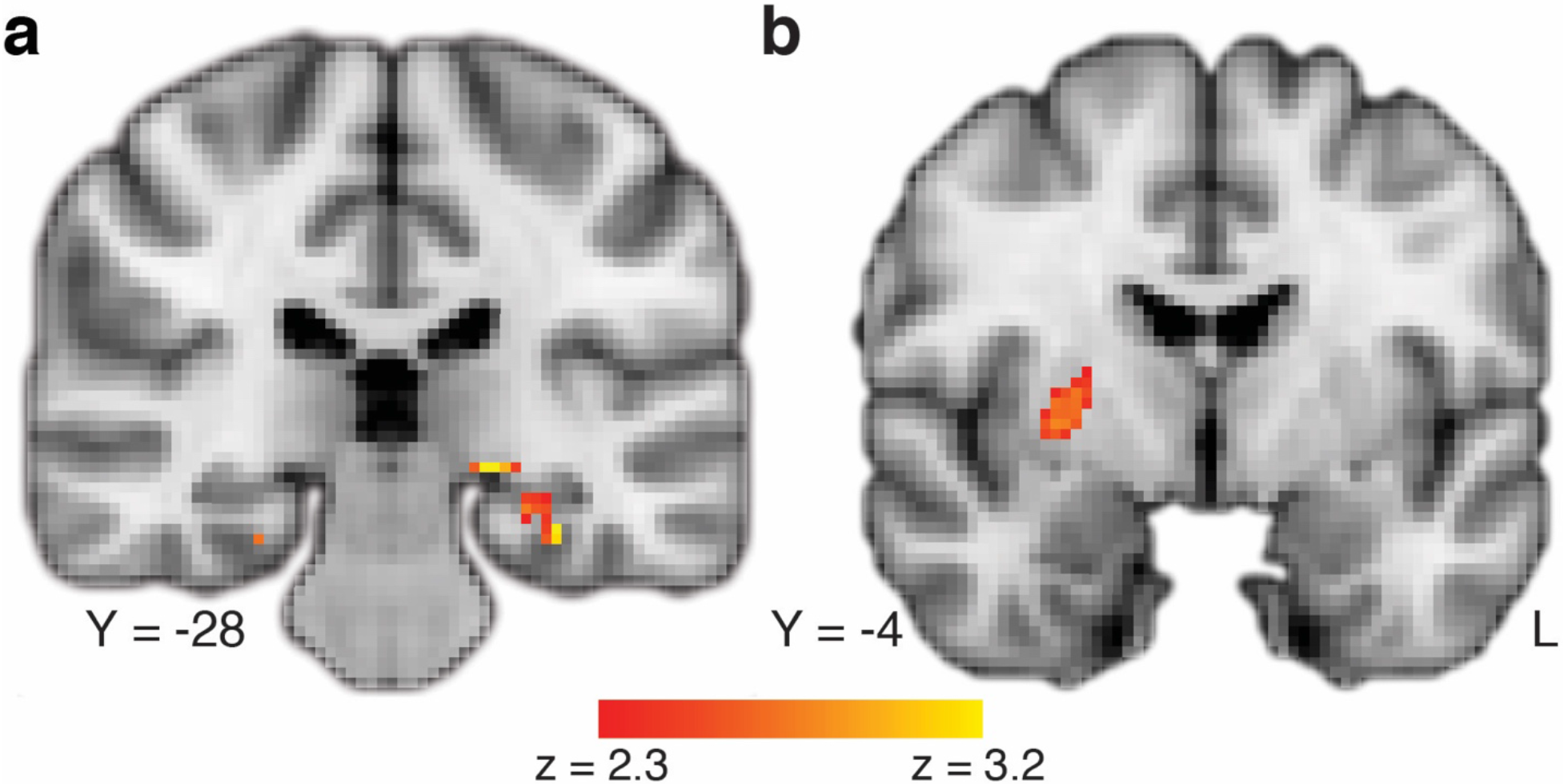
Timing of value-based decisions is related to activation in memory-localized regions of the hippocampus and to interactions between the hippocampus and the striatum. **a)** Areas of the brain that respond to memory retrieval are also more active with slower reaction times, especially for value-based decisions. This map exploits all three task conditions and shows a comparison of the effect of trial-by-trial RT on value-based decisions with perceptual decisions, localized (with a conjunction analysis) to regions of the brain that also show a memory-retrieval effect. The full map can be viewed at https://neurovault.org/collections/BOWMEEOR/images/56727. This effect in the hippocampus was replicated with an independent analysis using an anatomically defined region of the hippocampus even when accounting for several control variables (e.g. mean value across items in a pair, **Supplementary Figure 4d**). **b)** Connectivity related to the speed of value-based decisions. There was stronger correlation between the hippocampus and the striatum when value-based decisions were slower. The full map can be viewed at https://neurovault.org/collections/BOWMEEOR/images/56733. Coordinates reported in standard MNI space. Heatmap color bars range from z-stat = 2.3 to 3.2. Maps were cluster corrected for familywise error rate at a whole-brain level (**a**) and small volume corrected (SVC) in the striatum (b), p < 0.05.

We considered several alternative explanations of this result. First, one might argue that the value task makes more demands on memory, because it requires identifying objects (**Supplementary Figure 3a** and **Supplementary Table 4**). However, this explanation does not account for the correlation between hippocampal BOLD and reaction time, as all the objects are equally familiar, and therefore an account based on object identification demands would predict an overall difference between the two tasks regardless of timing of responses. In other words, there is no reason a priori to expect object identification demands to correlate with ΔValue or RT. Second, there is more variability in RT in the value-based task than in the perceptual task, but this alone does not explain our findings, as we find a similar result in the hippocampus when we conduct a repeated analysis restricting the RT to the range of overlap between the two tasks (**Supplementary Figure 3b** and **Supplementary Table 5**). Third, we explored whether overall difficulty could be contributing to these differences. Because RT is itself a function of the difficulty levels in the two tasks, we repeated the same analysis, controlling for the magnitude of color coherence and ΔValue, as well as other potential correlates of RT (e.g. mean of the pair of values; see *Methods*). This analysis again revealed RT-related activity in the hippocampus that is greater for the value-based than perceptual decisions (**Supplementary Figure 4** and **Supplementary Tables 6-8**). **Supplementary Figure 4d** shows the parameter estimates tracking trial-by-trial RT on BOLD, controlling for all these factors in a region of interest (ROI) independently defined by a contrast of successful memory retrieval in the hippocampus.

These results suggest a role for memory-related hippocampal mechanisms in accounting for value-based decisions. However, the hippocampus does not control decisions and actions directly. Thus, we were interested in exploring the broader neural circuits that interact with the hippocampus during value-based decisions and require longer deliberation times. We used a psychophysiological interaction (PPI) analysis to identify brain regions with activity that covaried in an RT-dependent manner with the activity of hippocampal “seed” voxels — i.e. those that exhibited RT-dependent activation on the value-based decision task and memory-related activation on the memory localizer task. The strongest RT-dependent correlation was between the hippocampus and the putamen, a region previously implicated in value learning and value-action coupling^29–32^. We previously hypothesized that the striatum would interact with the hippocampus for value-based decisions^15^ (**Figure 3b** and **Supplementary Table 9**). The PPI analysis tells us that the striatum functionally couples with the hippocampus in the service of value-based decisions that take more time to resolve, and while the analysis does not inform directionality, it suggests a possible mechanism by which value and memory associations can interact to guide action selection.

The findings suggest that decisions about value preference selectively recruit memory-related regions of the hippocampus proportionally to decision time. We reason that the brain seeks additional evidence when values associated with the pair of items are similar, perhaps evaluating the items along dimensions suggested by their juxtaposition (e.g., saltiness). Such evaluations might repeat sequentially until a clear preference is achieved. Although we cannot measure the content of participants’ thought process directly, we can test predictions that emerge from this framework. In particular, if the process of making a decision entails construction of value, then the items might undergo changes in value that affect subsequent encounters with the item, a process we refer to as “revaluation”.

To test this prediction, we exploited the design of our experiment, in which each of the 60 items was presented several times and compared to different items. We reasoned that if the decision process led to a change of value, then the next time that item is shown, its value might differ from its original value by an amount ±*δ*. The value would be incremented by if the item was chosen and reduced by this amount if it was rejected. Importantly, the algorithm we used to implement this idea is conservative: although we assume the process arises during the deliberation leading to a decision, we use only the outcome of the decision as an indicator of whether the revaluation was an increment or a decrement. In other words, updating only affects subsequent trials, not the ΔValue on the trial on which the revaluation takes place. We fit *δ* for each participant to minimize the deviance of a logistic choice function to the data.

A series of analyses confirmed that revaluation takes place, and that the outcome of our revaluation algorithm provides a better fit to the behavioral and brain data. For all participants, the difference in the revised values (Δ*V_rev_*) provided better fits of the choice and RT data, compared to the original values from the auction (**Figures 4a, Supplementary Figure 5** and **Supplementary Table 1**). The combined log likelihood improved by more than two orders of magnitude (**Figure 4b**, mean ΔBIC per participant: −53.4; p < 0.0001, Wilcoxon signed-rank test). Critically, such improvement is not guaranteed by the fitting procedure, and it is not observed in simulated data (**Supplementary Figure 6a**). It is also not unique to our dataset: We have validated the procedure using data from Folke et al.^33^ (**Supplementary Figure 6b**). Moreover, the logic behind the revaluation analysis predicts that the order of the trials should matter, because revaluation is assumed to be sequential and to follow the order of the choices that are made. We confirmed this by repeating the analysis on the same trials with the order permuted (**Supplementary Figure 6c**; p < 0.0001). Although our revaluation algorithm applies *V_rev_* to the next decision in which an item is encountered, our idea is that the construction of value occurs during the decision itself. In support of this, we found that the amount of revaluation (*δ*) was larger for short RT (**Supplementary Figure 6d**; p = 0.008, t-test), perhaps because deliberation led quickly to a compelling dimension of comparison to resolve preference. This result provides some evidence for a link between revaluation and the deliberation process itself.

**Figure 4:**
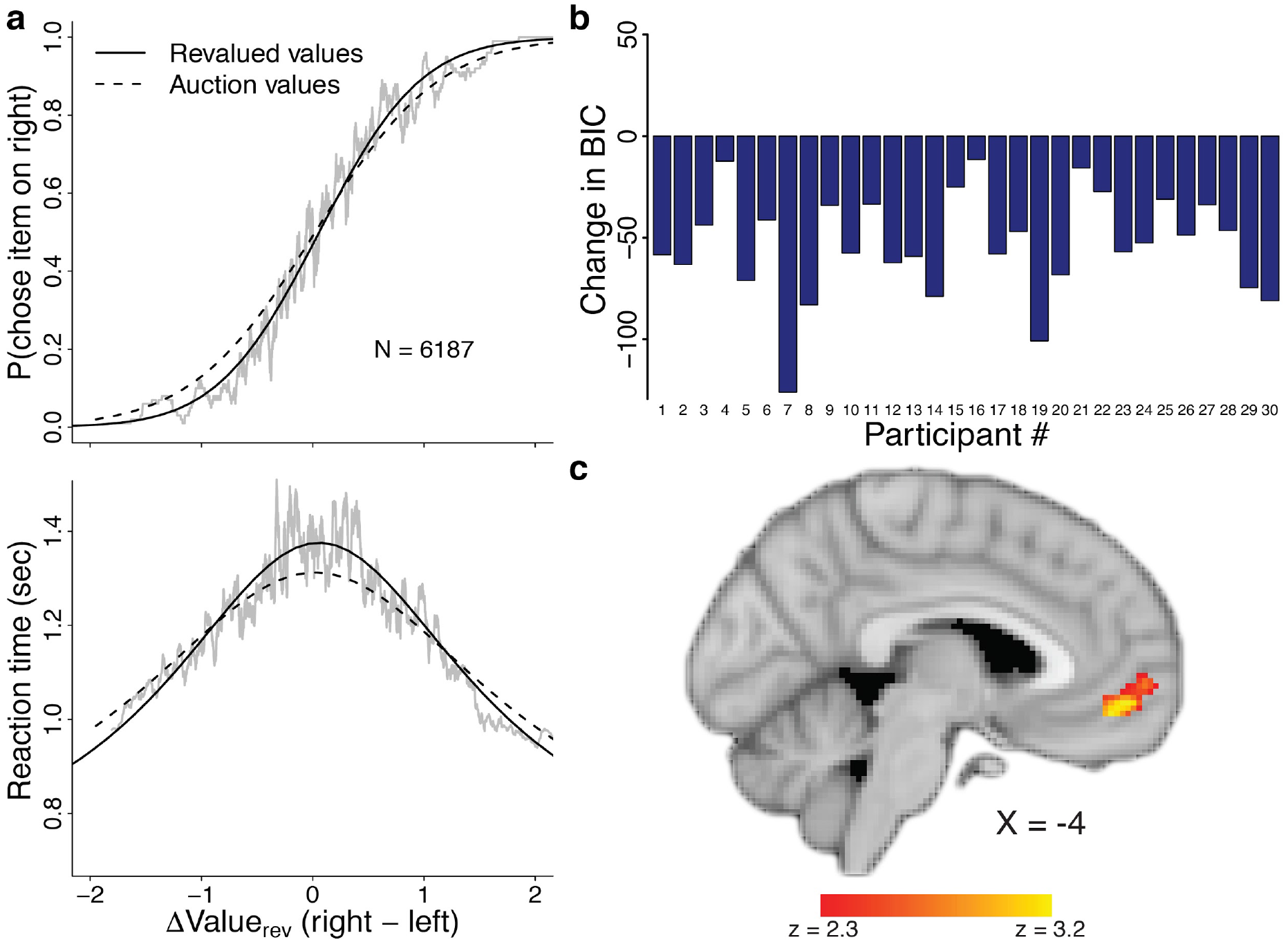
Revaluation of items occurs during decision making, affecting later choices, model fits, and value-related BOLD activity in ventromedial prefrontal cortex. **a)** Analysis of choices and RT using the revalued values shows improved fits to behavior. Gray lines are running means. Solid black lines are model fits based on revalued values. Dashed black lines are model fits based on original auction values. **b)** The change in Bayesian Information Criterion (BIC) for drift diffusion model fits based on the original auction values versus the fits based on revalued values per participant show that revaluation improved the behavioral fits for all participants, even when accounting for the additional parameter fit by the revaluation algorithm. **c)** The parametric effect of the revalued value of the chosen item shows a robust correlation between *V_rev_* and activity in the vmPFC. This model also includes the original auction value of the chosen item. To see the full map, go to https://neurovault.org/collections/BOWMEEOR/images/56734. Coordinates reported in standard MNI space. Heatmap color bars range from z-stat = 2.3 to 3.2. This map was cluster corrected for familywise error rate at a whole-brain level p < 0.05.

Additional support for a constructive revaluation process is adduced from the 60 pairs of items that were repeated during the experiment. Participants were more likely to exhibit an inconsistent preference if the items had been encountered more times between repetitions (**Supplementary Figure 6e**; p = 0.001), and they were also more likely to respond inconsistently if the Δ*V_rev_* diminished (odds ratio 5.3 relative to consistent; p < 0.0001, Fisher exact test). Finally, if revaluation renders a more accurate assessment of value, we predicted that the neural representation of value might reflect *V_rev_*. Whole-brain regression of BOLD activity against the *V_rev_* of the chosen item (controlling for auction value) revealed a robust response in the ventromedial PFC (**Figure 4c** and **Supplementary Table 10**), a region previously shown to respond to subjective value^34–37^.

Our findings demonstrate that time taken to indicate a preference between two food items is associated with activation in the hippocampus, particularly in regions that are also associated with memory retrieval, in the same participants. This RT-related activation was prominent in value-based decisions as compared to perceptual decisions that involved similar deliberation times and similar choice stochasticity. Our findings also show that specific items undergo revaluation, consistent with a constructive role for memory in guiding decisions. These findings thus support our hypothesis that the sequential samples of evidence bearing on preference arise through a process that makes use of memory, mediated in part by the hippocampus.

The findings do not tell us the exact role that the hippocampus plays, but it is almost certainly more nuanced than memory retrieval of the value associated with the items. Such associative memories are not thought to depend on the hippocampus^38–43^, and it is not obvious why a simple associative process would account for longer deliberation times or lead to changes in value. Instead, we favor the view that memory processes arise during decision making to construct value and to modify value along dimensions prompted by the comparison itself. The memory might not be about the items per se but play a role in the thought processes that guide evaluation and prospection^22,44^. Naturally, such deliberation is more likely to be evoked when items are valued similarly. Our analysis of revaluation supports a novel prediction of this theory: that the process of deciding could alter the value of the items themselves. We reason that this revaluation amounts to only a subtle revision because (i) the values derived from the preliminary auction are informative throughout the experiment and (ii) the best fitting update value, *δ*, is on the order of 1-10% of the original auction value. That said, we do not take the value of *δ* too seriously, because it is merely a proxy for a complex process.

One alternative explanation to account for the relation between choices and RT is that near-value choices take more time because they are based on noisy representations of value and these noisy quantities (or their differences) must be integrated to a threshold. This idea derives from the success of evidence accumulation models to reconcile accuracy and RT on a wide variety of cognitive and perceptual tasks^4,28,45,46^. In perceptual decision making using random dot motion, the neural representation of momentary evidence is clear as there are many representations of the accumulation of such evidence by neurons that represent the running accumulation of noisy samples^47,48^. A similar process is thought to underlie decisions in the dynamic random dot color dominance task used here^49^ and supported by **Figure 2** and **Supplementary Figure 7**. For value-based decisions, we suggest that instead the samples are derived from a constructive process involving memory. These samples might be combined (e.g., integrated), as is commonly assumed^4,50,51^, or the samples might be evaluated in turn until any one sample satisfies a criterion. This alternative retains consistency with the important insight that the process of deciding induces a revision in value itself.

Prior work has shown a role for the hippocampus in construction of value when making “new” flexible decisions^44,52,53^. Here we show a role for the hippocampus in guiding decisions about highly familiar items — decisions that need not, it seems, rely on flexible memory at all. The finding that memory plays a role in the construction of value during deliberation might have implications for other well-known phenomena in judgement and decision making, such as inconsistent preferences^54–56^, violations of transitivity^57^, and post-decision changes in valuation^58–60^, at least in some cases. In particular, it is worth noting that a memory-based account of value construction and revaluation offers an alternative explanation for one of the most extensively studied observations in judgement and decision making — that decision makers alter their value assessments after making a decision to justify previous choices, commonly referred to as “cognitive dissonance”^58^. We suggest that in the process of making a decision, items are evaluated and revalued and therefore what has been described as *post hoc* assessment to minimize dissonance may, in some cases, reflect a change in value made in service of the decision itself.

Whereas perceptual decisions are based mainly on objective evidence obtained from sources external to the organism, most of the decisions we make involve deliberation over evidence whose bearing on preference is fundamentally subjective. Of course, many decisions involve both data bearing on propositions as well as subjective preference, as in perceptual decisions by an artist or percipient. Thus, the idea that evidence is partially constructed through cognitive operations involving memory is likely to apply broadly, and it should not surprise us that it could lead to apparent inconsistency and revision. Value, like deliberation itself, evolves.

## Methods

### Participants

Thirty-three healthy participants were recruited through fliers posted on campus and surrounding area in New York City. Three participants were excluded from analysis due to excessive motion during MRI scanning. The final sample consisted of N = 30 (19 female), mean age = 24.7 ± 5.5 and self-reported Body Mass Index (BMI) = 23 ± 4.5. No statistical method was employed to pre-determine the sample size. The sample size we chose is similar to that used in previous publications.

All experimental procedures were approved by the Institutional Review Board (IRB) at Columbia University and all participants provided signed informed consent before taking part in the study.

### Tasks

The study took place over two sessions. On the first day, participants were not scanned. They were trained on the perceptual color dots task (details below), received feedback (correct/incorrect) on each trial during training, and were trained to criterion, defined as 80% or higher accuracy over the last four blocks of 10 trials. Training consisted of a minimum 200 trials and a maximum 400 trials. After color dots training, participants underwent incidental encoding for the Memory Localizer task: they rated 100 neutral objects, presented one at a time on the computer screen, on how much they liked that object by placing the cursor along a visual analog scale that ranged from 0 (least) to 10 (most) using the computer mouse. This liking rating task served as a memory encoding phase, followed two days later by a surprise memory recognition test in the scanner (details below). The first study session lasted about one hour. When it ended, participants were told to refrain from eating or drinking anything besides water for four hours before their next appointment. On the second session, which took place two days after the first session, participants took part in an auction outside of the scanner. They were then placed in the MRI scanner and performed the food choice task, the color dots task, and the memory recognition task.

#### Auction

Participants were endowed with $3, which they used to take part in an auction. The auction followed Bercker-Degroot-Marschak (BDM) rules^61^. This auction procedure allowed us to obtain a measure of willingness-to-pay (WTP) for each of 60 appetitive food items per participant^35^. Participants were presented with one snack item at a time, in random order, on a computer screen. They placed their bid by moving a cursor on an analog scale that spanned from $0 to $3 at the bottom of the screen using the computer mouse. The auction was self-paced, and the next item was presented only after the participant placed their bid. After participants placed bids on all 60 items, they were given a chance to revise their bids to account for adjustments and scaling effects that can occur after participants experienced the full food stimulus set. Participants were presented with each of the 60 items in random order a second time with their original bid displayed below and were asked whether they wanted to change their bid. If they clicked “NO”, they were presented with the next food item, and their original bid was kept as the WTP for that item. If the participant clicked “YES”, the $0 to $3 analog scale was presented and they placed a new bid using the mouse as before. This new bid was recorded as the final WTP for that item. The starting location of the cursor along the analog scale was randomized on each trial and the mouse cursor was reset to the middle of the screen on each trial to prevent participants from simply clicking through the entire auction phase without deliberation. Participants were told that a single trial would be drawn at random at the end of the session, and that they could bid any amount of the full $3 for each food item and would not spread their endowment over multiple items. They were told that their best strategy to win the auction was to bid exactly what each item was worth to them to purchase from the experimenter at the end of the experiment and that bidding consistently high or consistently low was a bad strategy. At the end of the session, the computer generated a counter bid in the form of a random number between $0 and $3 in increments of 25 cents. If the computer bid was higher than the participant’s bid, then he or she lost the auction; if the participant matched or outbid the computer, they were offered to purchase the randomly drawn food item from the experimenter at the computer’s bid lower price. The outcome of the auction was played out at the end of the experimental session. After performing the auction outside the scanner, participants performed the following three tasks in the scanner while functional brain images were acquired.

#### Food Choice

The 60 food items were rank-ordered based on WTPs obtained during the auction, and 150 unique pairs of items were formed such that the difference in WTP between the two items in a pair (i.e. ΔValue) varied. Pairs were presented in random order, one pair at a time, with one item on each side of a central fixation cross. Right/left placement of the higher value item in a pair was counterbalanced across trials. Participants were instructed to choose the food they preferred. Participants chose one item on each trial by pressing one of two buttons on an MRI-compatible button box. They were given up to 3 s to make their selection. After a choice was made, the selected item was highlighted for 500 ms. If participants did not make a choice before the 3 s cutoff, the message “Please respond faster” was displayed for 500 ms. Trials were separated by a jittered inter trial interval (ITI) drawn from a truncated exponential distribution with a minimum 1 s and a maximum 12 s and a mean of 3 s. Participants were told that they would be given the chosen food on a single randomly selected trial to eat. Participants were presented with 210 trials total, split into three 70-trial scan runs. Runs of the food choice were interleaved with runs of the color dots task (below). Of the 150 unique pairs, 90 pairs were presented only once and 60 pairs were presented twice. Thus, each of the 60 food items was presented 7 times in total. Each scan run of the food choice task lasted 7 min.

#### Color Dots

Participants viewed a dynamic display with flickering dots and were asked to determine whether there were more yellow or blue dots. Participants responded by pressing one of two buttons, with the color-button mapping counterbalanced across participants. The visual stimulus consisted of many dots that were either yellow or blue. On each trial, the stimulus was assigned a color coherence that determined the dominant color category and its ambiguity. The sign of the color coherence determined which color category the stimulus belonged to: positive meant blue and negative meant yellow. The absolute magnitude of the color coherence determined ambiguity of the color category: the smaller the absolute coherence, the more ambiguous the category. Formally, the color coherence is a log odds ratio of each dot being yellow (versus blue) on each frame, i.e., *log*(*p*_*yellow*_/(1−*p*_*yellow*_)). Note that although *p*_*yellow*_ is fixed within a trial, the actual number of yellow dots in each frame will fluctuate frame by frame. The color coherence was drawn from a set of 11 values (−2, −1, −0.5, −0.25, −0.125, 0, 0.125, 0.25 0.5, 1, 2) on each trial. All dots jumped to a random position within a circular aperture (5° diameter) on every frame. The stimulus was presented for a maximum of 2.5 s. Participants were instructed to make their response as soon as they had an answer. If they responded within the 2.5 s window, the stimulus disappeared and a central fixation cross would appear. During scanning on session 2, participants did not receive feedback on whether they responded correctly, unlike during training on session 1 where they received correct/incorrect feedback on each trial. Feedback on session 1 appeared after a response was made and remained on the screen for 500 ms. If they did not respond within the 2.5 s choice window, a message asking the participant to please respond faster was displayed for 500 ms. Trials were separated by jittered ITIs that were drawn from a truncated exponential distribution with a minimum of 1 s, maximum of 12 s and a mean of 3 s. Participants were presented with of a total of 210 trials, split into three scan runs of 70 trials each. Each scan run of the color dots task lasted 6 min and 30 s.

#### Memory Recognition

Participants were presented with the 100 objects that they rated during session 1 as well as 100 new objects, randomly intermixed, one object at a time in the middle of the screen. Below the image of the object and to the right and left of center appeared the words “OLD” and “NEW” that corresponded to the right/left button mapping. On each trial, participants were asked to determine whether the object on the screen was old, meaning they remember rating that object on their first visit or if the object is new and they do not remember seeing or rating that object. Participants responded by pressing one of two buttons on an MRI-compatible button box. Old/New response-button mapping was counterbalanced across participants. The stimulus remained on the screen for a maximum of 3 s. If participants responded within the 3 s response window, their choice (i.e. OLD or NEW) was highlighted for 500 ms. If they did not respond within the 3 s window, a message asking them to please respond faster was displayed for 500 ms. Trials were separated by a jittered ITI drawn from a truncated exponential distribution (min 1 s, max 12 s, mean 3 s). The 200 trials were split into four scan runs of 50 trials (approximately 5 min) each. All four runs of this task were consecutive, with no intervening other tasks in between.

### fMRI acquisition

Imaging data were acquired on a 3 T GE MR750 MRI scanner with a 32-channel head coil. Functional data were acquired using a T2*-weighted echo planar imaging sequence (repetition time (TR) = 2 s, echo time (TE) = 22 ms, flip angle (FA) = 70°, field of view (FOV) = 192 mm, acquisition matrix of 96 × 96). Forty oblique axial slices were acquired with a 2 mm in-plane resolution positioned along the anterior commissure-posterior commissure line and spaced 3 mm to achieve full brain coverage. Slices were acquired in an interleaved fashion. Each of the food choice runs consisted of 212 volumes, each of the color dots runs consisted of 197 volumes, and the memory test runs consisted of 150 volumes. In addition to functional data, a single three-dimensional high-resolution (1 mm isotropic) T1-weighted full-brain image was acquired using a BRAVO pulse sequence for brain masking and image registration.

### Behavioral analysis

#### Choice and reaction time data

Choice and RT data were analyzed using regression models. Choice data were scored on accuracy (consistency of responses with the stated value or WTP for the choice option, i.e. score 1 for trials when the participant chose the food with higher WTP and 0 if they chose the food with lower WTP). These binary data were then entered into a repeated measures logistic regression mixed effects model to calculate the odds of choosing consistently with their prior valuation and its relationship to |ΔValue|. RT data were entered into a repeated measures linear regression mixed model to test the relationship between RT and |ΔValue|.

#### Drift Diffusion model

We fit a one-dimensional drift-diffusion model to the choice and RT on each decision. The model assumes that choice and reaction time are linked by a common mechanism of evidence accumulation, which stops when a decision variable reaches one of two bounds. The decision variable (*x*) is given by the cumulative sum of samples from a Normal distribution with mean *μdt* and variance *dt*,

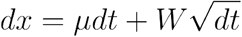

where *W* is the Wiener process. The accumulation starts with x = 0.

In the value-based decision, the mean of the momentary evidence is given by

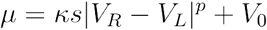

where *V_R_* and *V_L_* are the values of the right and left item respectively, *k* is the signal-to-noise, and *V_0_* is a bias to account for non-symmetric distributions of choice or RT between left and right choices. We allowed the drift-rate to be a nonlinear function of ΔValue, assuming a power law transformation with parameter. represents the sign of the value difference (positive if *V_R_* > *V_R_*
, negative otherwise). Similarly, in the color-discrimination task the mean of the momentary evidence is given by

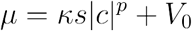

where *c* is the color coherence, and *s* is positive if *c* > 0 and negative otherwise. We used time-varying decision bounds to account for potential differences in RT between correct and error trials. The shape of the bound was determined by three parameters. The bound is flat (with height *B_0_*) from time zero to time *B_d_*, and then collapses exponentially towards zero with time constant *B_2_* (in seconds). The two bounds were assumed to be symmetrical around x = 0. For the value-based task, the positive bound represents a commitment to a rightward choice, and the negative bound represents a commitment to a leftward choice. For the perceptual task, the positive and negative bounds indicate a commitment to the blue and yellow choice, respectively. The reaction time is given by the sum of the decision time, determined by the drift-diffusion process, and a non-decision time that we assume Gaussian with mean *t_nd_* and standard deviation *σ_tnd_*.

We performed three different set of fits: (i) perceptual decision task, (ii) value-based task with original BDM auction values, and (iii) value-based task with values from revaluation (see below). The model was fit to maximize the joint likelihood of choice and RT of each trial. The likelihood of the parameters given the data from each trial was obtained by numerically solving the Fokker-Planck equations describing the dynamics of the drift-diffusion process. For the value-based task, we reduced the number of unique drift-rates by rounding ΔValue to multiples of 0.1 dollar. In **Figure 2**, we fit the model to grouped data from all participants. The fits for individual participants are shown in **Supplementary Figures 1** and **5**. The best fitting parameters for the grouped and non-grouped data are shown in **Supplementary Table 1**.

#### Revaluation algorithm

Each of the 60 food items was presented seven times. Following the order of trials in the experiment, we incremented or decremented the value of each food item by *δ* depending on whether the item was chosen or unchosen, respectively. The items were assigned their auction value on first encounter, and the updated values, *V_rev_*, only affected the estimate of ΔValue (used to fit choice and RT) on subsequent trials. We fit *δ* for each participant to minimize the deviance of a logistic choice function to the data. Although fitting *δ* must reduce the deviance, on average, but it does not guarantee that *δ* would be positive or at all substantial. To test this, we ran the algorithm on simulated choices derived by sampling from the logistic fits to each participant’s choice data (**Supplementary Figure 6a**). To evaluate the importance of trial order, we applied the revaluation algorithm to 1000 random permutations of each participant’s trials (**Supplementary Figure 6 b** & **c**). To test if the amount of revaluation depends on the duration of the deliberation process leading to a choice, we fit two *δ* per participant, using short and long RTs based on a median split (**Supplementary Figure 6d**).

### Imaging analysis

#### Imaging data preprocessing

Raw imaging data in DICOM format were converted to NIFTI format and preprocessed through a standard preprocessing pipeline using the FSL package version 5^62^. Functional image time series were first aligned using the MCFLIRT tool to obtain six motion parameters that correspond to the x-y-z translation and rotation of the brain over time. Next, the skull was removed from the T2* images using the brain extraction tool (BET) and from the high-resolution T1 images using Freesurfer^63,64^. Spatial smoothing was performed using a Gaussian kernel with a full-width half maximum (FWHM) of 5 mm. Data and design matrix were high-pass filtered using a Gaussian-weighted least-squares straight line fit with a cutoff period of 100 s. Grand-mean intensity normalization of each run’s entire four-dimensional data set by a single multiplicative factor was performed. The functional volumes for each participant and run were registered to the high resolution T1-weighted structural volume using a boundary-based registration method implemented in FSL5 (BBR^BBR, 65^). The T1-weighted image was then registered to the MNI152 2 mm template using a linear registration implemented in FLIRT (12 degrees of freedom). These two registration steps were concatenated to obtain a functional-to-standard space registration matrix.

#### Food choice

We conducted a GLM analysis on the food choice task data. The first analysis was the simplest and included three regressors of interest: (i) onsets for all valid choice trials; (ii) same onsets and duration as (i) but modulated by RT; (iii) onsets for missed trials. After running this model, we ran a conjunction analysis using the output of this model and the equivalent model on the perceptual decision task data (see below) with our main memory retrieval success contrast (see memory recognition section below). The conjunction map is presented in **Figure 3a**. This model was also used to generate the map in **Supplementary Figure 3a**.

The second GLM analysis was designed to rule out the possibility that differences in RT variance between the two tasks might account for a contrast between tasks in the effect of RT on BOLD. This model included five regressors of interest: (i) onsets for all valid choice trials with RT in the range of overlap across the two tasks; (ii) same onsets and duration as (i) but modulated by RT; (iii) onsets for all valid choice trials with RT not in the range of overlap across the two tasks; (iv) onsets for missed trials. This model was used to generate the map in **Supplementary Figure 3b**.

The GLM model we based our inferences on included twelve regressors of interest: (i) onsets for non-repeated unique pair “correct” trials (i.e. unique pairs of items that were only presented once where choice was consistent with initial valuation during the auction meaning the chosen item had the higher WTP), modeled with a duration that equaled the average RT across all valid food choice trials and participants; (ii) same onsets and duration as (i) but modulated by |ΔValue| demeaned across these trials within each run for each participant; (iii) same onsets and duration as (i) but modulated by RT demeaned across these trials within each run for each participant; (iv-vi) similar to regressors (i-iii), but for non-repeated unique pair “incorrect” trials (i.e. unique pairs of items that were only presented once for which choice was inconsistent with initial valuation during the auction, meaning the chosen item had the lower WTP); (vii-ix) similar to regressors (i-iii), but for repeated unique pair trials (i.e. unique pairs of items that were presented twice, both “correct” and “incorrect” trials together); (x) to account for any differences in mean value across items in a pair (i.e. average WTP across both items in a pair) between trial types, we added a regressor with the onsets of all valid trials and the same duration as all other regressors, while the modulator was the demeaned mean WTP across both items in a pair; (xi) to account for any differences in right/left choices between trial types, we added a regressor with the same onsets and durations as (x), while the modulator was an indicator for right/left response; (xii) onsets for missed trials. Parameter estimates from this model are presented in **Supplementary Figure 4d** and a map from this model is presented in **Supplementary Figure 4a**.

Finally, we conducted a GLM analysis to look at BOLD activity related to revalued values (from the revaluation algorithm described above). This model included eight regressors of interest. (i) onsets for all valid trials, modeled with a duration which equaled the average RT across all valid food choice trials and participants; (ii) same onsets and duration as (i) but modulated by |ΔValue_auction_| demeaned across these trials within each run for each participant; (iii) same onsets and duration as (i) but modulated by |ΔValue_rev_| demeaned across these trials within DVARS, each run for each participant; (iv) same onsets and duration as (i) but modulated by RT demeaned across these trials within each run for each participant; (v) same onsets and duration as (i) but modulated by V_auction_ of the chosen item demeaned across these trials within each run for each participant; (vi) same onsets and duration as (i) but modulated by V_rev_ of the chosen item demeaned across these trials within each run for each participant; (vii) to account for any differences in right/left choices between trial types we added a regressor with the same onsets and durations as (i), while the modulator was an indicator for right/left response; (viii) onsets for missed trials. The map in **Figure 4c** was generated using this model.

In all models, we also included the six x, y, z translation and rotation motion parameters obtained from MCFLIRT, framewise displacement (FD) and RMS intensity difference from one volume to the next^DVARS, 66^ as confound regressors. We also modeled out volumes with FD and DVARS that exceeded a threshold of 0.5 by adding a single time point regressor for each “to-be-scrubbed” volume ^67^. All regressors were entered at the first level of analysis and all (but the added confound regressors) were convolved with a canonical double-gamma hemodynamic response function. The temporal derivative of each regressor (except for the added confound regressors) was included in the model. The models were estimated separately for each participant and each run.

#### Color dots

The first GLM analysis was the simplest and included three regressors of interest: (i) onsets for all valid choice trials; (ii) same onsets and duration as (i) but modulated by RT; (iii) onsets for missed trials. After running this model, we ran a conjunction analysis using the output of this model and the equivalent model on the value-based decision task data (see above) with our main memory retrieval success contrast (see memory recognition section below). The conjunction map is presented in **Figure 3a**. This model was also used to generate the map in **Supplementary Figure 3a**.

The second GLM analysis was run to rule out the possibility that differences in RT variance between the two tasks might account for a contrast between tasks in the effect of RT on BOLD. This model included five regressors of interest: (i) onsets for all valid choice trials with RT in the range of overlap across the two tasks; (ii) same onsets and duration as (i) but modulated by RT; (iii) onsets for all valid choice trials with RT not in the range of overlap across the two tasks; (iv) onsets for missed trials. This model was used to generate the map in **Supplementary Figure 3b**. The GLM model for the color dots task that we based our inferences on included 3 regressors for each of correct and incorrect color choice trial types: (i) onsets of correct trials (i.e. participant chose yellow when the coherence was negative and chose blue when the coherence was positive, as well as all coherence 0 trials) modeled with a duration which equaled the average RT across all valid color dots trials and participants; (ii) same onsets and durations as (i) but modulated by |color coherence| demeaned across these trials within each run for each participant; (iii) same onsets and durations as (i) but modulated by RT demeaned across these trials within each run for each participant; (iv-vi) similar to regressors (i-iii), but for incorrect trials (i.e. participant chose yellow when the coherence was positive and chose blue when the coherence was negative); additionally we included two other regressors (vii) to account for any differences in right/left choices between trial types we added a regressor with the onsets of all valid color dots trials and the same duration as all other regressors (average RT across all trials and participants), while the modulator was an indicator for right/left response; (viii) onsets for missed trials. Parameter estimated from this model are presented in **Supplementary Figure 4d**, and a map from this model is presented in **Supplementary Figure 4b**.

For all models, we added the same covariates as in the food choice design matrix, including the six motion regressors described above, along with FD and DVARS as confound regressors.

#### Memory recognition

The GLM for the memory recognition task data included 8 regressors of interest: (i) onsets of hit trials (i.e. participant responded old when the object on the screen was old), modeled with a duration which equaled the average RT across all valid memory trials and participants; (ii) same onset and duration as (i) but modulated by liking rating for the object demeaned across these trials within each run for each participant; (iii) onsets of miss trials (i.e. participant responded new when the object on the screen was old) modeled with the same duration as (i); (iv) same onset and duration as (iii) but modulated by liking rating for the object demeaned across these trials within each run for each participant; (v) onsets of correct rejection trials (i.e. participant responded new when the object on the screen was new) modeled with the same duration as (i); (vi) onsets of false alarm trials (i.e. participant responded old when the object on the screen was new) modeled with the same duration as (i); (vii) to account for any differences in RT between trial types we added a regressor with the onsets of all valid trials and the same duration as all other regressors (average RT across all trials and participants) while the modulator was the demeaned RT across all valid trials; (viii) onsets for missed trials. We added the same covariates as in the food choice design matrix, including the six motion regressors described above, along with FD and DVARS as confound regressors. The map for the contrast hits > correct rejections in this model is presented in **Supplementary Figure 2**. This contrast was also used in the conjunction analysis presented in **Figure 3a**.

#### Conjunction analysis

To test the spatial overlap in memory-retrieval-related brain activity and value-based RT related activation, we conducted a conjunction analysis between the maps presented in **Supplementary Figure 2** (memory retrieval success contrast of hits [regressor (i) in memory recognition fMRI GLM model] greater than correct rejections [regressor (v) in memory recognition fMRI GLM model]) and the same contrast as in **Supplementary Figure 4c**, but for the simpler model (contrast of value-based RT [regressor (iii) in the first food choice fMRI GLM model] greater than perceptual RT [regressor (iii) in the first color dots fMRI GLM model]). The conjunction map is presented in **Figure 3a**.

#### Psychophysiological interaction (PPI)

As the seed for the PPI analysis, we used significant voxels for the contrast value-based RT greater than perceptual RT (**Supplementary Figure 4c**) that lay within an anatomical mask of bilateral hippocampus (Harvard-Oxford Atlas). PPI regressors were created by deconvolving the seed to obtain an estimated neural signal during value-based decisions using SPM’s deconvolution algorithm^68^, calculating the interaction with the task in the neural domain and then reconvolving to create the final regressor. We followed McLaren et al.’s gPPI modeling procedure^69^ and included 9 regressors in our GLM: (i) onsets of lower-than-median RT trials, modeled with a duration which equaled the average RT across all valid trials and participants; (ii) onsets of higher-than-median RT trials, modeled with the same duration as in (i); (iii) onsets of all valid trials, modeled with the same duration as in (i) and modulated by |ΔValue|, demeaned across these trials within each run for each participant; (iv) same onsets and duration as (iii) but modulated by the average value across the two paired foods, demeaned across these trials within each run for each participant; (v) to account for any differences in right/left choices, we added a regressor with the same onsets and duration as (iii) but modulated by an indicator for right/left response; (vi) onsets of all missed trials with the same duration as (i); (vii) the raw time course extracted from the seed (after registering the seed to the native space of each run for each participant); (viii) a PPI regressor with the same onsets as (i); (ix) a PPI regressor with the same onsets as (ii). The PPI between lower and higher RT food choice trials generated the map in **Figure 3b**.

## Data and code availability

Data from this study are available from the corresponding author upon request. Task code and analysis code is available at https://github.com/abakkour/MDMRT_scan. Imaging analysis code is available from the corresponding author upon request.

## Acknowledgements

This research was funded by the McKnight Memory and Cognitive Disorders Award (D.S.), National Science Foundation Directorate for Social, Behavioral & Economic Sciences Postdoctoral Research Fellowship grant #1606916 (A.B.), National Institutes of Health grant R01EY011378 and Howard Hughes Medical Institute (M.N.S.). The authors would like to thank Yul HR Kang for developing the Color Dots task as well as helpful discussions, Daniel Kimmel, Danique Jeurissen, and David Barack for feedback on an earlier draft of the manuscript and helpful discussions. The authors would also like to thank Lucy Owen, Sean Raymond, Eileen Hartnett, and the New York State Psychiatric Institute MR Unit staff for help with data collection.

## Author contributions

A.B., M.N.S. and D.S. designed the experiment. A.B. coded the experiment. A.B. conducted the experiment. A.B. and A.Z. analyzed the data. A.B., A.Z., M.N.S. and D.S. discussed the results and wrote the paper.

## Additional information

Correspondence and request for materials should be addressed to A.B.

## Supplementary information for Value-based decisions involve sequential sampling from memory

### Supplementary Text

#### Behavioral Results

##### Value-based Decisions

Participants tended to choose the item that was of higher value (as measured by willingness-to-pay in the auction phase), and this tendency increased as the value difference between the two items (i.e. ΔValue) increased (Figure 2A top, the odds ratio of choosing the item on the right side of the screen as ΔValue increased was 17.19, 95% CI = [10.97 26.93], p < 0.0001). Participants’ reaction times increased as |ΔValue| decreased (Figure 2A, bottom); β = −0.13, bootstrapped 95% CI [-0.18 −0.08], bootstrapped p = 0.001). These results replicate previous findings that show that choices and reaction times vary systematically with ΔValue (*6*, *70*). These relationships are captured by the drift-diffusion model (solid black lines in Figure 2), suggesting that the mechanism underlying the decision is based on accumulation of evidence.

##### Perceptual Decisions

When performing the Color Dots task, participants responded blue more often as the color coherence increased and responded yellow more often as the color coherence decreased (became more negative, Figure 2B top, the odds ratio of choosing blue-to-yellow as color coherence increased was 120.31, 95% CI [64.23 225.38], p < 0.0001). RTs were shortest at the lowest and highest color coherence levels. RTs were longest at color coherence level zero, when there was an equal proportion of yellow and blue dots in the stimulus (Figure 2B bottom). In a repeated measures linear regression mixed effects model, color coherence was negatively related to RT (β = −0.24, bootstrapped 95% CI [-0.27 −0.20], bootstrapped p = 0.001).

##### Memory recognition

Participants’ mean hit rate was 0.81 ± 0.15, and their mean correct rejection (CR) rate was 0.76 ± 0.15. Mean *d’* across all participants was 1.78 ± 0.58. Participants were faster when making a correct response (hits and CRs combined, mean RT = 1.23 ± 0.17) than when making an incorrect response (misses and false alarms combined, mean RT = 1.39 ± 0.24, mean of the differences in RT = 0.16, 95% CI [0.12 0.22], t(29) = 7.21, p < 0.0001). The liking ratings for objects on session 1 were related to responses and RTs during the memory recognition test on session 2. In a repeated measures logistic regression mixed effects model including only data for old objects seen on session 1, the odds of responding old-to-new as the liking rating increased was 1.12 (95% CI [1.03 1.22], p = 0.011). In a repeated measures linear regression mixed effects model, liking rating was weakly negatively related to RT (β = −0.006, bootstrapped 95% CI [-0.012 −0.001], bootstrapped p = 0.02).

### Supplementary Figures

**Supplementary Figure 1:**
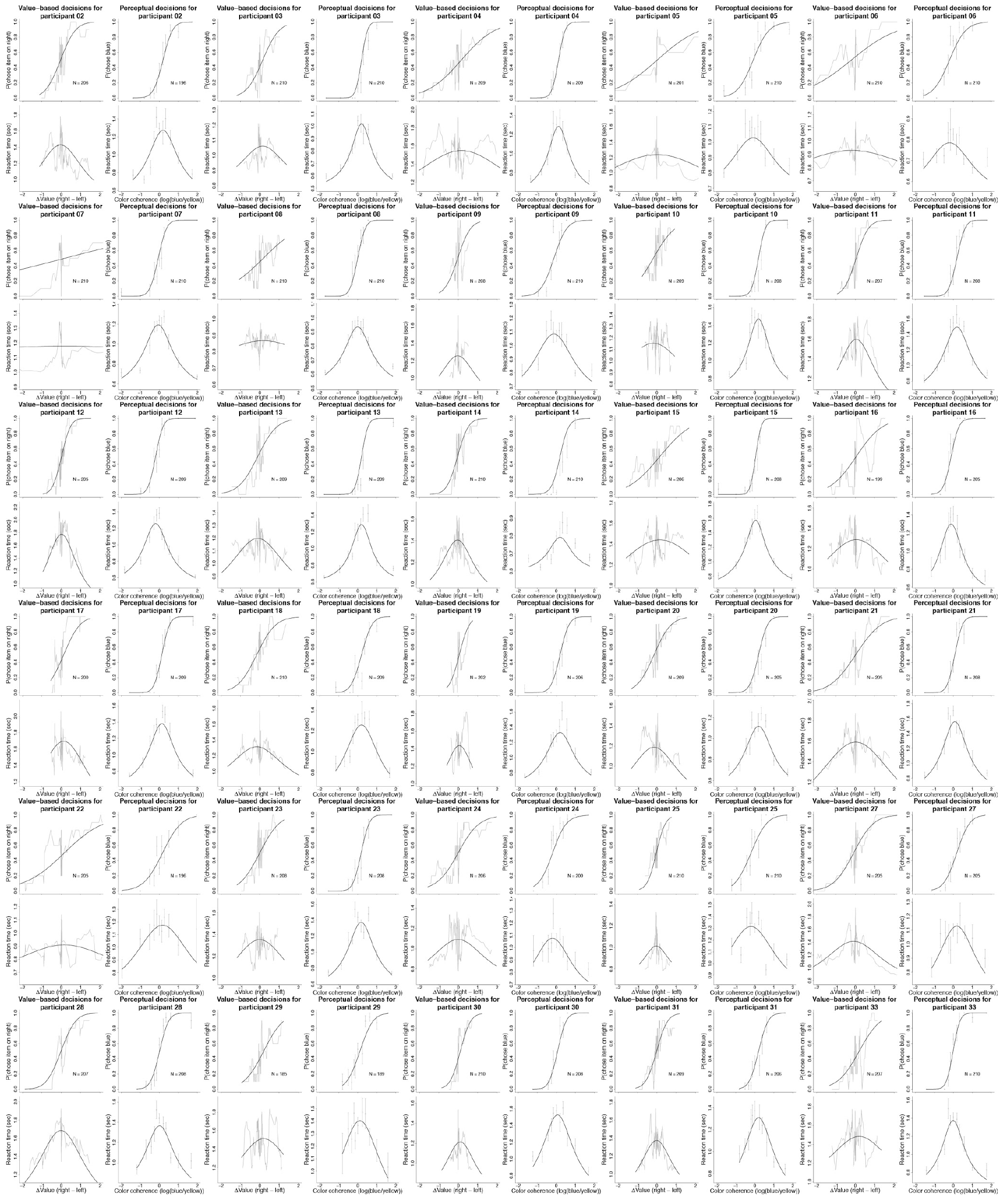
Data and DDM fits for value-based and perceptual decisions per participant. Gray lines are a running means. Gray dots are means and grey error bars are standard errors of the mean. Solid black lines are model fits.

**Supplementary Figure 2:**
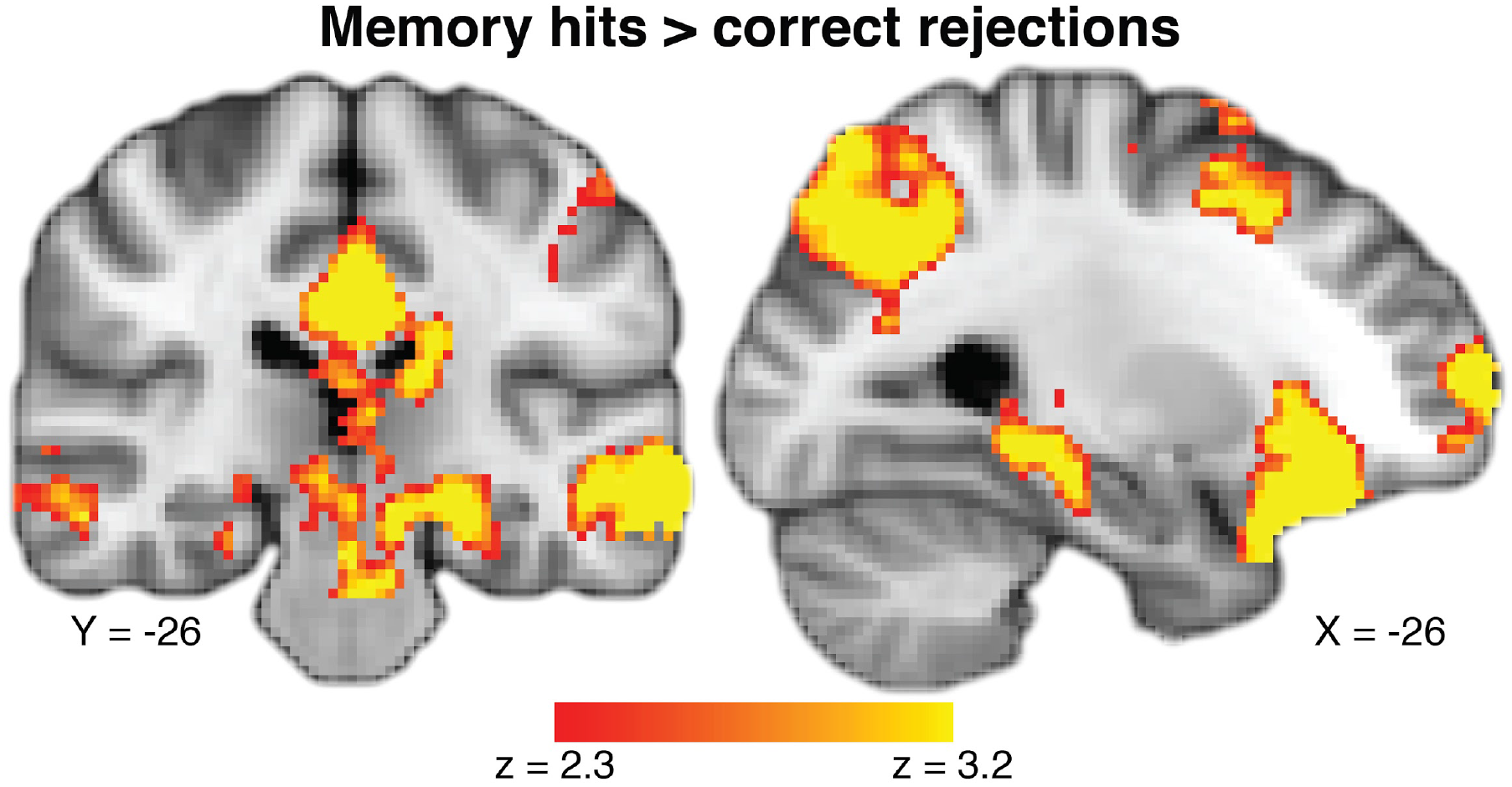
Parametric map of main effect of hits versus correct rejections during memory recognition. This map was used in the conjunction map presented in **Figure 3a**. Coordinates reported in standard MNI space. Heatmap color bars range from z-stat = 2.3 to 3.2. These maps were cluster-corrected at a whole-brain level p < 0.05. To see the full uncorrected map, go to https://neurovault.org/collections/BOWMEEOR/images/56726.

**Supplementary Figure 3:**
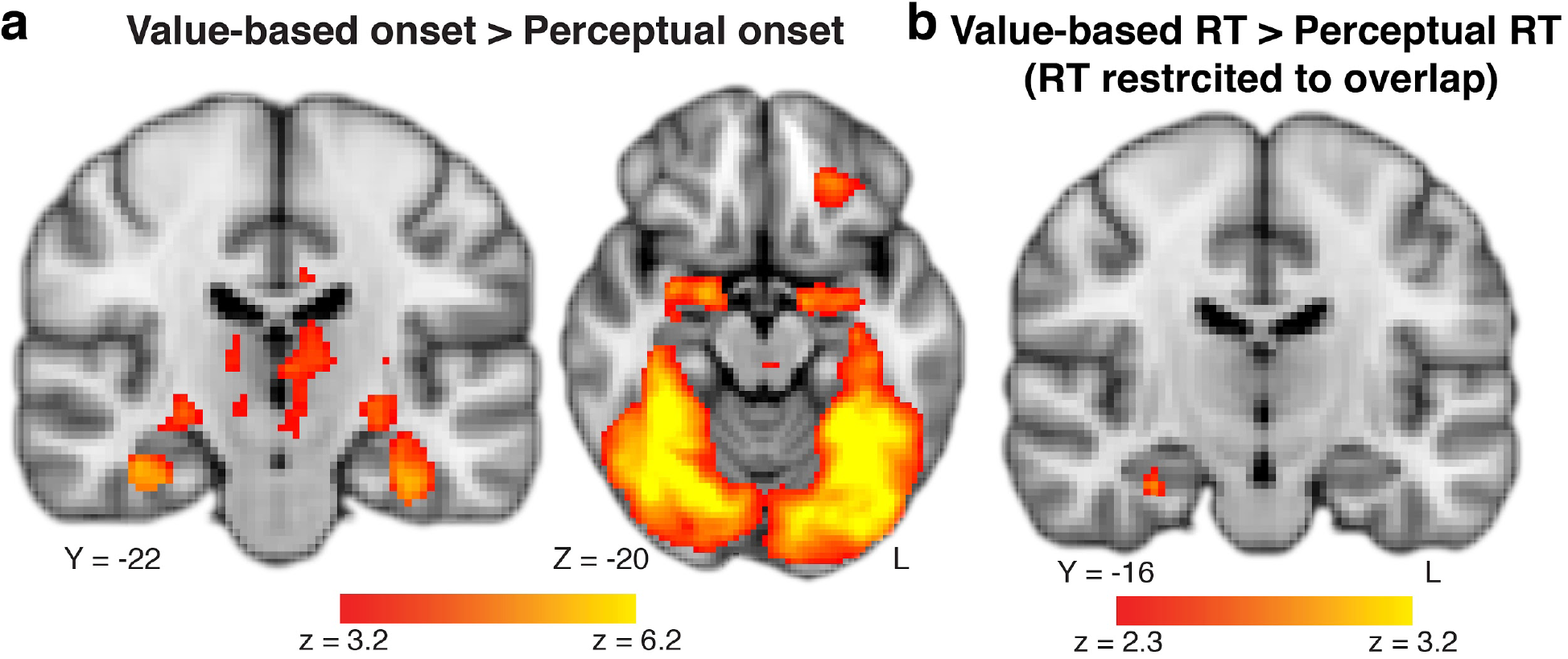
**a)**Main effect of task type on whole-brain BOLD activity at stimulus onset. We find very strong ventral stream and hippocampus activation for the value-based compared to the perceptual decision task. This is not surprising as ventral stream activity is crucial to recognition of objects such as the food items used. To see the full uncorrected map, go to https://neurovault.org/collections/BOWMEEOR/images/56728**. b)** Modulated effect of value-based compared to perceptual RT on whole-brain BOLD activity, restricted to the range of RTs that overlap across the two tasks. Even when restricting RT variance to be equivalent across value-based and perceptual decision tasks, we see a more positive relationship between BOLD in the hippocampus and RT during value-based when compared to perceptual decisions, confirming the results from a simpler model in **Figure 3a** and a more complex model in **Supplementary Figure 4c&d**. To see the full uncorrected map, go to https://neurovault.org/collections/BOWMEEOR/images/56729. Coordinates reported in standard MNI space. Heatmap color bars range from z-stat = 3.2 to 6.2 in (**a**) and z-stat = 2.3 to 3.2 in (**b**). These maps were cluster-corrected at a whole-brain level p < 0.05.

**Supplementary Figure 4:**
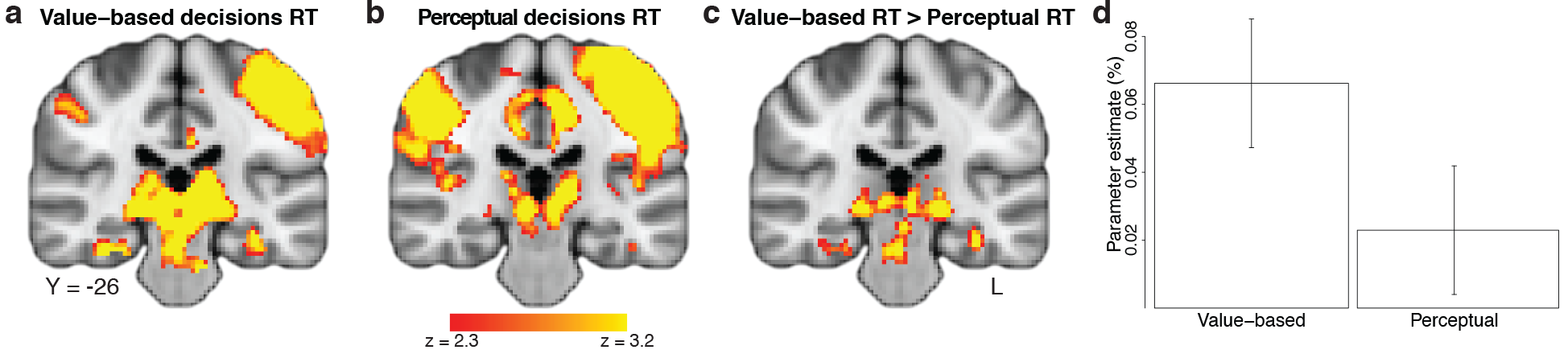
Parametric maps of the modulated effect of reaction time on BOLD in **a)** value-based (to see the full uncorrected map, go to https://neurovault.org/collections/BOWMEEOR/images/56731) and **b)** perceptual decisions (to see the full map, go to https://neurovault.org/collections/BOWMEEOR/images/56732) separately in a model that includes several regressors (e.g. mean of pair values, see *Methods*.) **c)** The contrast between maps in (**a**) and (**b**) (to see the full map, go to https://neurovault.org/collections/BOWMEEOR/images/56730). **d)** Mean parameter estimates for the effect of RT on BOLD in value-based and perceptual decisions from an independent ROI defined by successful memory retrieval activity (**Supplementary Figure 2**) within an anatomical mask of the hippocampus. Coordinates reported in standard MNI space. Heatmap color bars range from z-stat = 2.3 to 3.2. These maps were cluster-corrected at a whole-brain level p < 0.05.

**Supplementary Figure 5:**
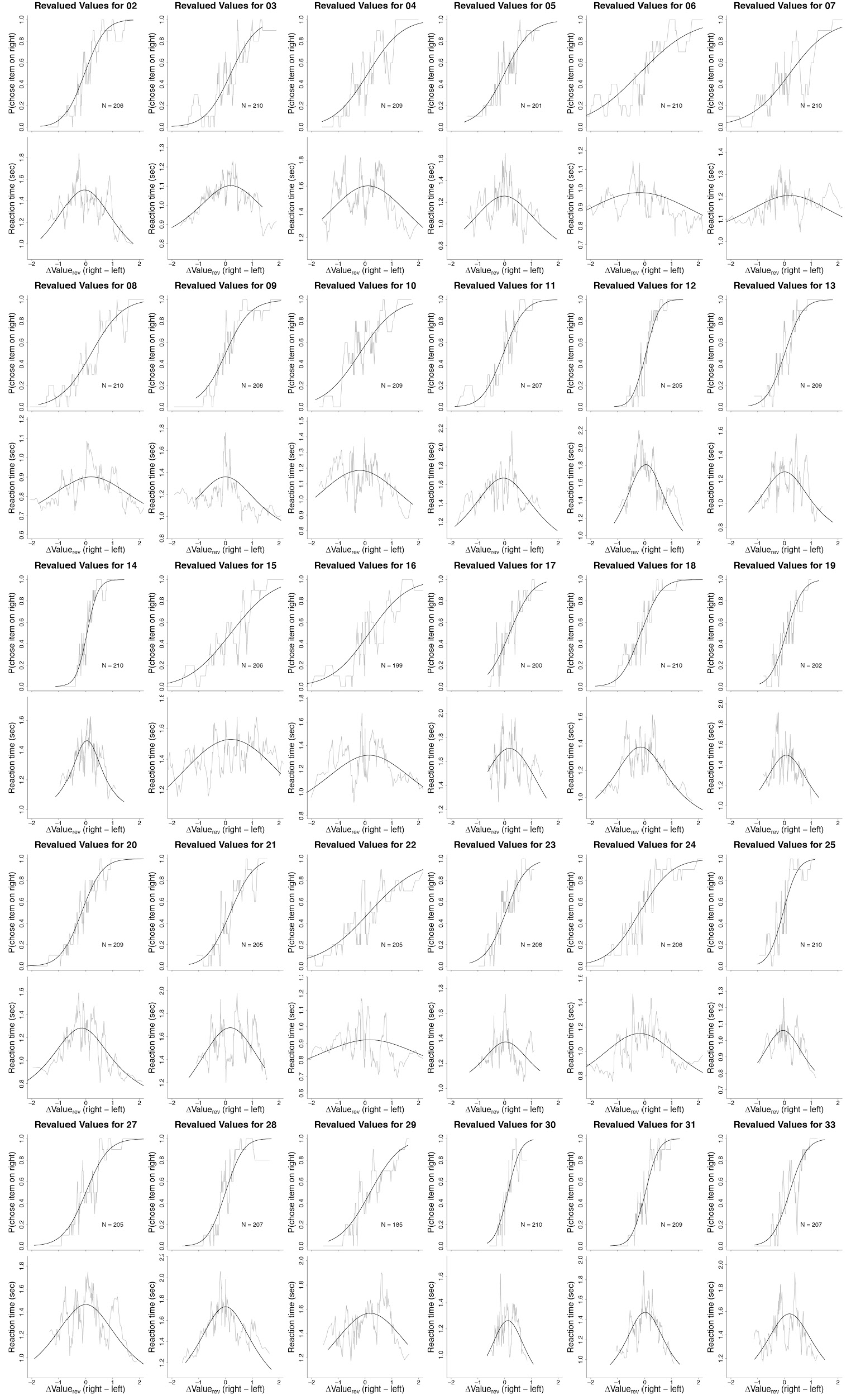
Reanalysis of behavior for each participant based on V_rev_. Gray lines are a running means. Solid black lines are model fits.

**Supplementary Figure 6:**
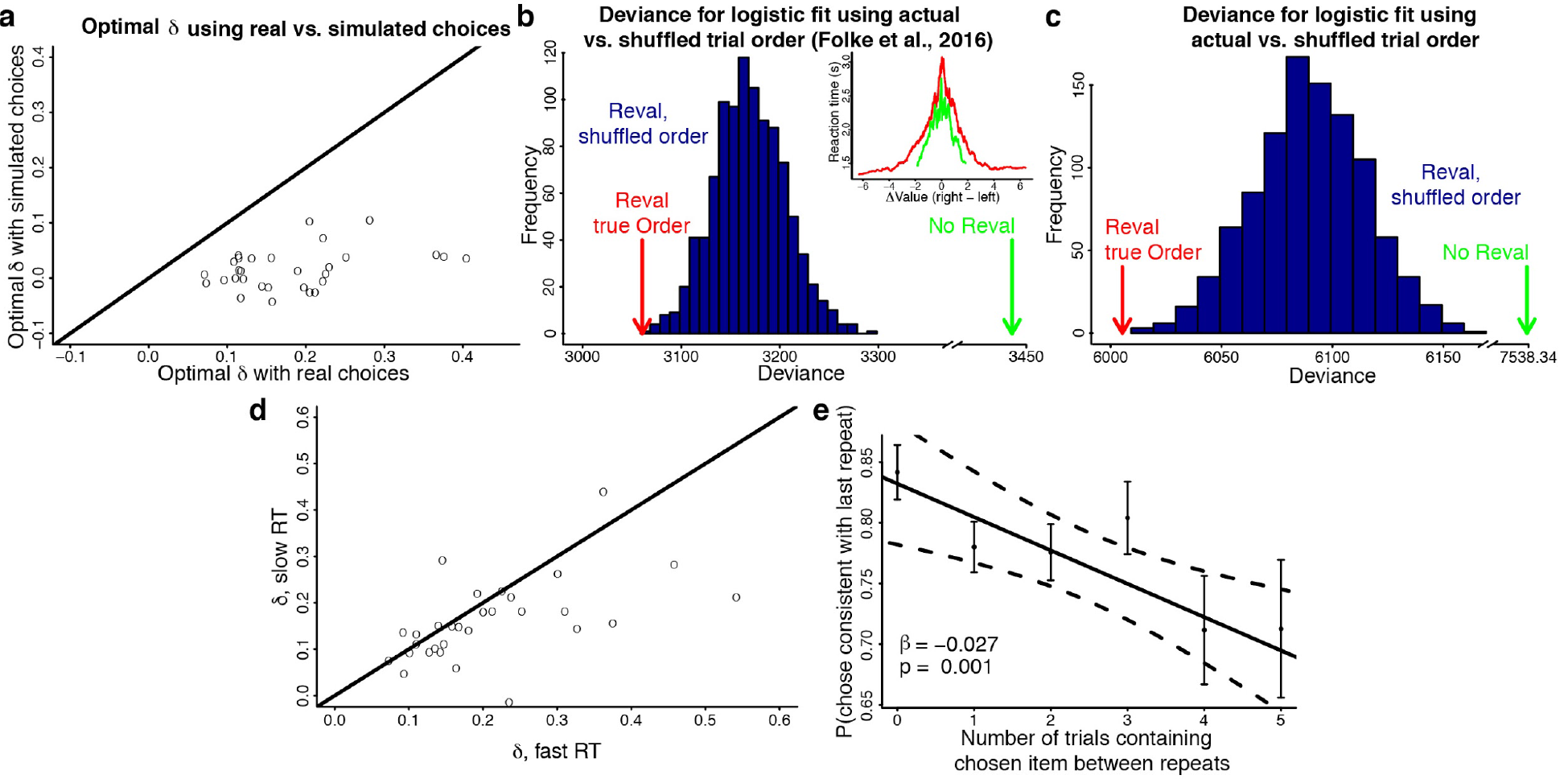
Additional analyses bearing on revaluation. **a)** Scatter plot compares (*i*) the revaluation update (*δ*) derived from each participant’s choices with (*ii*) the *δ* derived from simulated stochastic choices based on the participant’s choice function (logistic fit). The *δ* estimated from the real data are greater than zero and they are greater than the derived from simulated data. **b, c)** Trial order dependence of revaluation in two data sets (**b**, Folke et al., 2016; **c**, our data set). Frequency histograms show the distribution of deviance (error) in the logistic fits of revalued data under random permutations of trial order. The true order leads to the best fit (red arrow). Revaluation of permuted order is better than the fit using auction derived values (green arrow). This is not surprising because all revaluations exploit a degree of freedom. The inset in (**b**) shows the running mean of reaction times across all trials and participants in the Folke et al. data set using the original values (green) and the revalued values (red). **d)** The magnitude of revaluation update (*δ*) depends on reaction time. Scatter plot shows *δ* for short and long RT (median split) for each participant. The updates tended to be larger when the RT was faster (p < 0.01, paired t-test). **e)** Repetitions of identical pairs of items lead to less consistent preference if the items have been considered in other trials. The graph shows the probability of a consistent choice as a function of the number of times the item in that pair that was chosen on the first repetition had appeared in other comparisons before the pair repeated. The number of times the unchosen item appeared affects consistency similarly (not shown). These item counts mediate the effect of time and number of trials. Combined data from 30 participants (N = 1726 trials); error bars are s.e.m. The solid black line is the regression line and dashed curved lines are 95% confidence intervals for the regression fit.

**Supplementary Figure 7:**
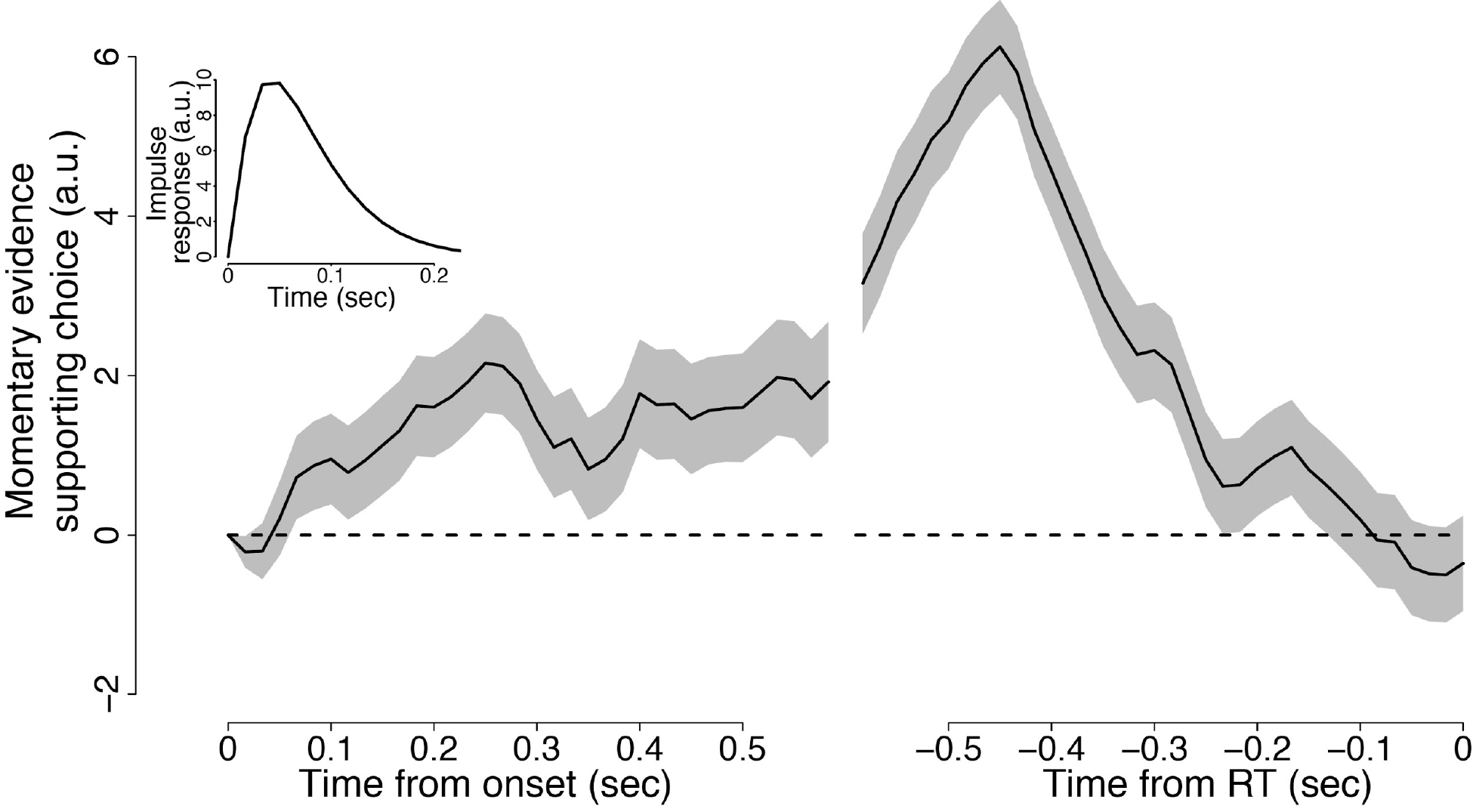
Effect of stimulus information on choice. The graphs depict the average excess of colored dots at each time point in support of the choice (i.e. blue minus yellow on trials in which the participant chose blue, and vice versa). The counts are derived from each video frame, and smoothed by the kernel (shown in the inset) and detrended by subtracting the mean for each color coherence condition. (**Left**) Stimulus data are aligned to onset. Only trials with RT>740 ms are included, and averages exclude stimulus data within 500 ms of the RT. It shows that information acquired at the beginning of the trial affect choices rendering later, consistent with evidence integration. (**Right**) Stimulus data are aligned to RT. Averages exclude stimulus data within 200 ms of onset. It shows that stimulus information ceases to affect the choice 300-400 ms, consistent with non-decision time inferred from model fits. Shading is s.e.m. Only color coherences 0 and ±0.125 are included in the analysis.

### Supplementary Tables

**Supplementary Table 1.**
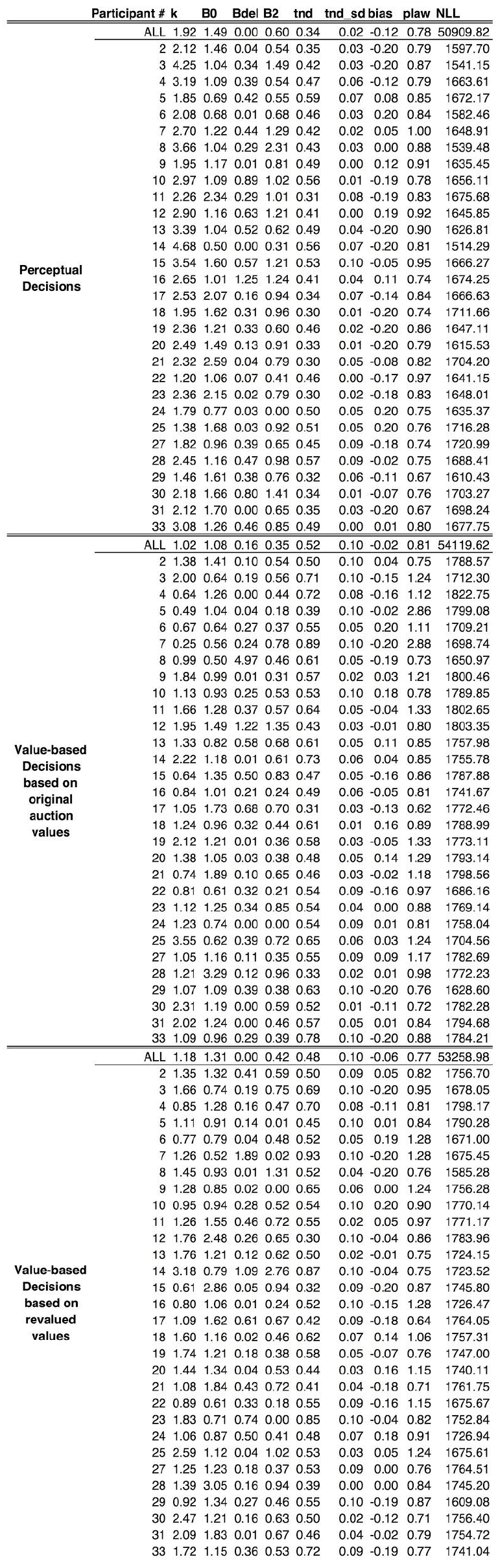
DDM parameter estimates.

**Supplementary Table 2.**
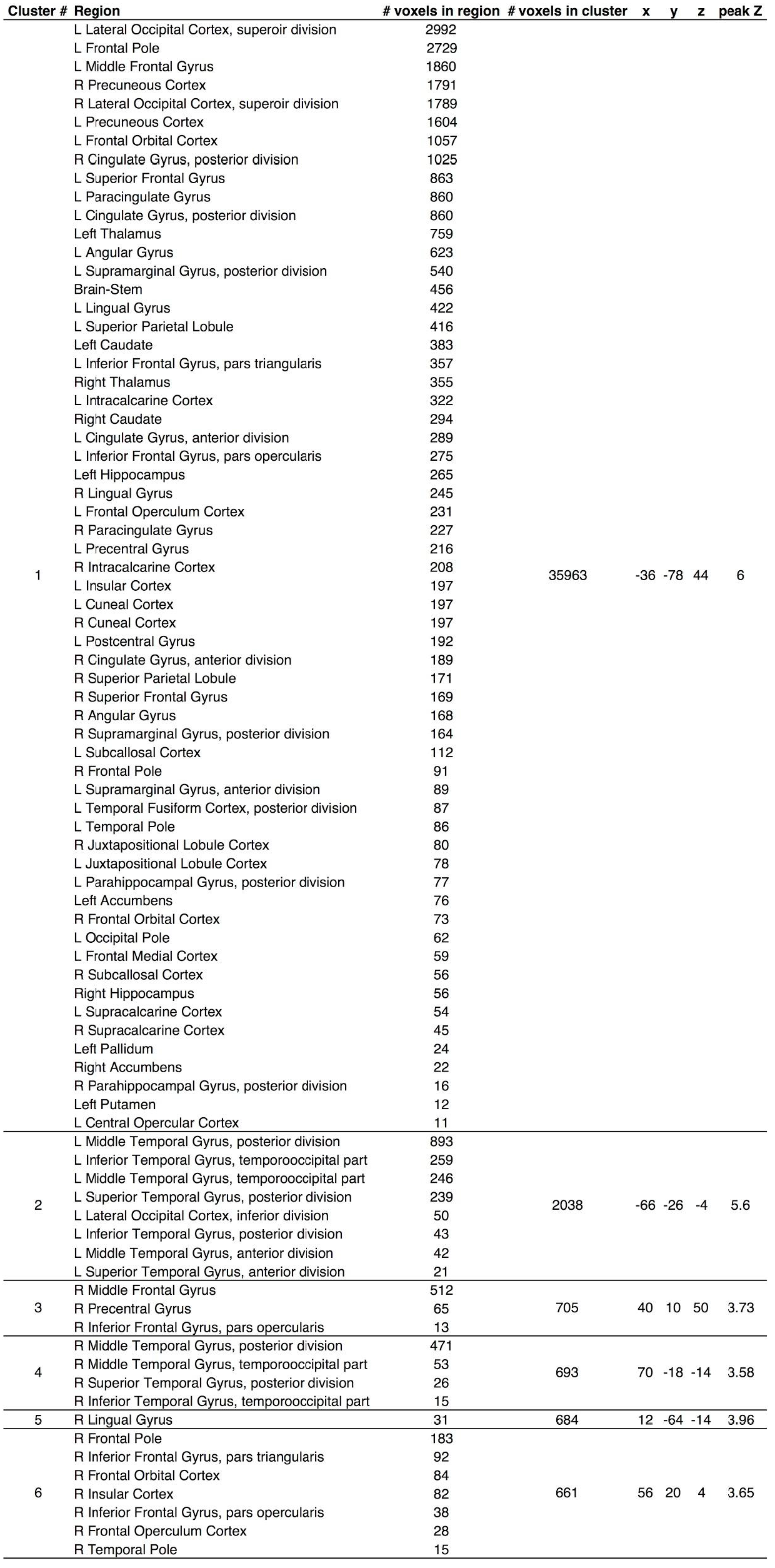
Activation table for map in Supplementary Figure 2. (successful memory retrieval: hits > correct rejections)

**Supplementary Table 3.** Activation table for map in Figure 3a. (conjunction between RT effect on BOLD for value-based greater than perceptual with effect of successful memory recognition)

| Cluster # | Region | # voxels in region | # voxels in cluster | x | y | z | peak Z |
| --- | --- | --- | --- | --- | --- | --- | --- |
| 1 | L Middle Frontal Gyrus | 345 |  |  |  |  |  |
|  | L Inferior Frontal Gyrus, pars triangularis | 93 | 514 | -42 | 30 | 10 | 4.08 |
|  | L Inferior Frontal Gyrus, pars opercularis | 11 |  |  |  |  |  |
| 2 | R Precuneous Cortex | 260 |  |  |  |  |  |
|  | R Intracalcarine Cortex | 29 | 382 | 20 | -66 | 24 | 4.61 |
|  | R Supracalcarine Cortex | 19 |  |  |  |  |  |
|  | R Cuneal Cortex | 16 |  |  |  |  |  |
| 3 | L Precuneous Cortex | 169 |  |  |  |  |  |
|  | L Intracalcarine Cortex | 65 | 347 | -14 | -68 | 12 | 4.75 |
|  | L Supracalcarine Cortex | 32 |  |  |  |  |  |
|  | L Cuneal Cortex | 27 |  |  |  |  |  |
| 4 | L Intracalcarine Cortex | 73 | 172 | -8 | -82 | 4 | 3.5 |
|  | L Lingual Gyrus | 56 |  |  |  |  |  |
| 5 | L Temporal Fusiform Cortex, posterior division | 63 | 89 | -34 | -42 | -22 | 3.72 |
|  | Left Hippocampus | 14 |  |  |  |  |  |
| 6 | Left Hippocampus | 29 | 68 | -18 | -32 | -2 | 4.22 |
|  | Left Thalamus | 19 |  |  |  |  |  |

**Supplementary Table 4.** Activation table for map in Supplementary Figure 3a. (overall main effect of value-based greater than perceptual decisions)

| Cluster # | Region | # voxels in region | # voxels in cluster | x | y | z | peak Z |
| --- | --- | --- | --- | --- | --- | --- | --- |
| 1 | R Lateral Occipital Cortex, superior division | 3635 | 53545 | 10 | -80 | 4 | 7.96 |
|  | L Lateral Occipital Cortex, superior division | 3277 |  |  |  |  |  |
|  | R Lingual Gyrus | 1777 |  |  |  |  |  |
|  | R Occipital Pole | 1765 |  |  |  |  |  |
|  | L Middle Frontal Gyrus | 1660 |  |  |  |  |  |
|  | L Occipital Pole | 1640 |  |  |  |  |  |
|  | L Lingual Gyrus | 1405 |  |  |  |  |  |
|  | R Precuneous Cortex | 1322 |  |  |  |  |  |
|  | L Lateral Occipital Cortex, inferior division | 1268 |  |  |  |  |  |
|  | R Lateral Occipital Cortex, inferior division | 1211 |  |  |  |  |  |
|  | L Precentral Gyrus | 1013 |  |  |  |  |  |
|  | L Occipital Fusiform Gyrus | 926 |  |  |  |  |  |
|  | R Occipital Fusiform Gyrus | 880 |  |  |  |  |  |
|  | Left Thalamus | 820 |  |  |  |  |  |
|  | R Temporal Occipital Fusiform Cortex | 796 |  |  |  |  |  |
|  | R Intracalcarine Cortex | 770 |  |  |  |  |  |
|  | R Cuneal Cortex | 725 |  |  |  |  |  |
|  | L Superior Frontal Gyrus | 710 |  |  |  |  |  |
|  | L Precuneous Cortex | 680 |  |  |  |  |  |
|  | L Temporal Occipital Fusiform Cortex | 648 |  |  |  |  |  |
|  | L Intracalcarine Cortex | 631 |  |  |  |  |  |
|  | Left Hippocampus | 619 |  |  |  |  |  |
|  | Right Thalamus | 612 |  |  |  |  |  |
|  | L Frontal Pole | 580 |  |  |  |  |  |
|  | L Superior Parietal Lobule | 568 |  |  |  |  |  |
|  | L Temporal Fusiform Cortex, posterior division | 545 |  |  |  |  |  |
|  | L Inferior Temporal Gyrus, temporooccipital part | 489 |  |  |  |  |  |
|  | L Cuneal Cortex | 486 |  |  |  |  |  |
|  | Brain-Stem | 466 |  |  |  |  |  |
|  | Right Hippocampus | 463 |  |  |  |  |  |
|  | R Cingulate Gyrus, posterior division | 403 |  |  |  |  |  |
|  | R Inferior Temporal Gyrus, temporooccipital part | 354 |  |  |  |  |  |
|  | L Cingulate Gyrus, posterior division | 304 |  |  |  |  |  |
|  | L Paracingulate Gyrus | 291 |  |  |  |  |  |
|  | R Temporal Fusiform Cortex, posterior division | 284 |  |  |  |  |  |
|  | Left Amygdala | 277 |  |  |  |  |  |
|  | L Frontal Orbital Cortex | 250 |  |  |  |  |  |
|  | R Superior Parietal Lobule | 248 |  |  |  |  |  |
|  | Right Amygdala | 244 |  |  |  |  |  |
|  | L Postcentral Gyrus | 212 |  |  |  |  |  |
|  | Left Putamen | 193 |  |  |  |  |  |
|  | R Supracalcarine Cortex | 189 |  |  |  |  |  |
|  | L Inferior Frontal Gyrus, pars triangularis | 188 |  |  |  |  |  |
|  | L Juxtapositional Lobule Cortex | 178 |  |  |  |  |  |
|  | L Inferior Frontal Gyrus, pars opercularis | 174 |  |  |  |  |  |
|  | R Parahippocampal Gyrus, anterior division | 162 |  |  |  |  |  |
|  | L Supramarginal Gyrus, anterior division | 149 |  |  |  |  |  |
|  | L Insular Cortex | 139 |  |  |  |  |  |
|  | R Paracingulate Gyrus | 135 |  |  |  |  |  |
|  | L Parahippocampal Gyrus, anterior division | 128 |  |  |  |  |  |
|  | L Parahippocampal Gyrus, posterior division | 128 |  |  |  |  |  |
|  | R Superior Frontal Gyrus | 115 |  |  |  |  |  |
|  | R Parahippocampal Gyrus, posterior division | 115 |  |  |  |  |  |
|  | L Cingulate Gyrus, anterior division | 87 |  |  |  |  |  |
|  | R Juxtapositional Lobule Cortex | 77 |  |  |  |  |  |
|  | R Cingulate Gyrus, anterior division | 74 |  |  |  |  |  |
|  | L Supracalcarine Cortex | 71 |  |  |  |  |  |
|  | L Central Opercular Cortex | 68 |  |  |  |  |  |
|  | R Angular Gyrus | 60 |  |  |  |  |  |
|  | L Supramarginal Gyrus, posterior division | 59 |  |  |  |  |  |
|  | L Inferior Temporal Gyrus, posterior division | 58 |  |  |  |  |  |
|  | L Angular Gyrus | 40 |  |  |  |  |  |
|  | Left Pallidum | 34 |  |  |  |  |  |
|  | Right Putamen | 19 |  |  |  |  |  |
| 2 | R Middle Frontal Gyrus | 1430 | 2506 | 36 | -6 | 46 | 4.29 |
|  | R Precentral Gyrus | 285 |  |  |  |  |  |
|  | R Inferior Frontal Gyrus, pars triangularis | 44 |  |  |  |  |  |
|  | R Frontal Pole | 32 |  |  |  |  |  |
|  | R Superior Frontal Gyrus | 15 |  |  |  |  |  |

**Supplementary Table 5.**
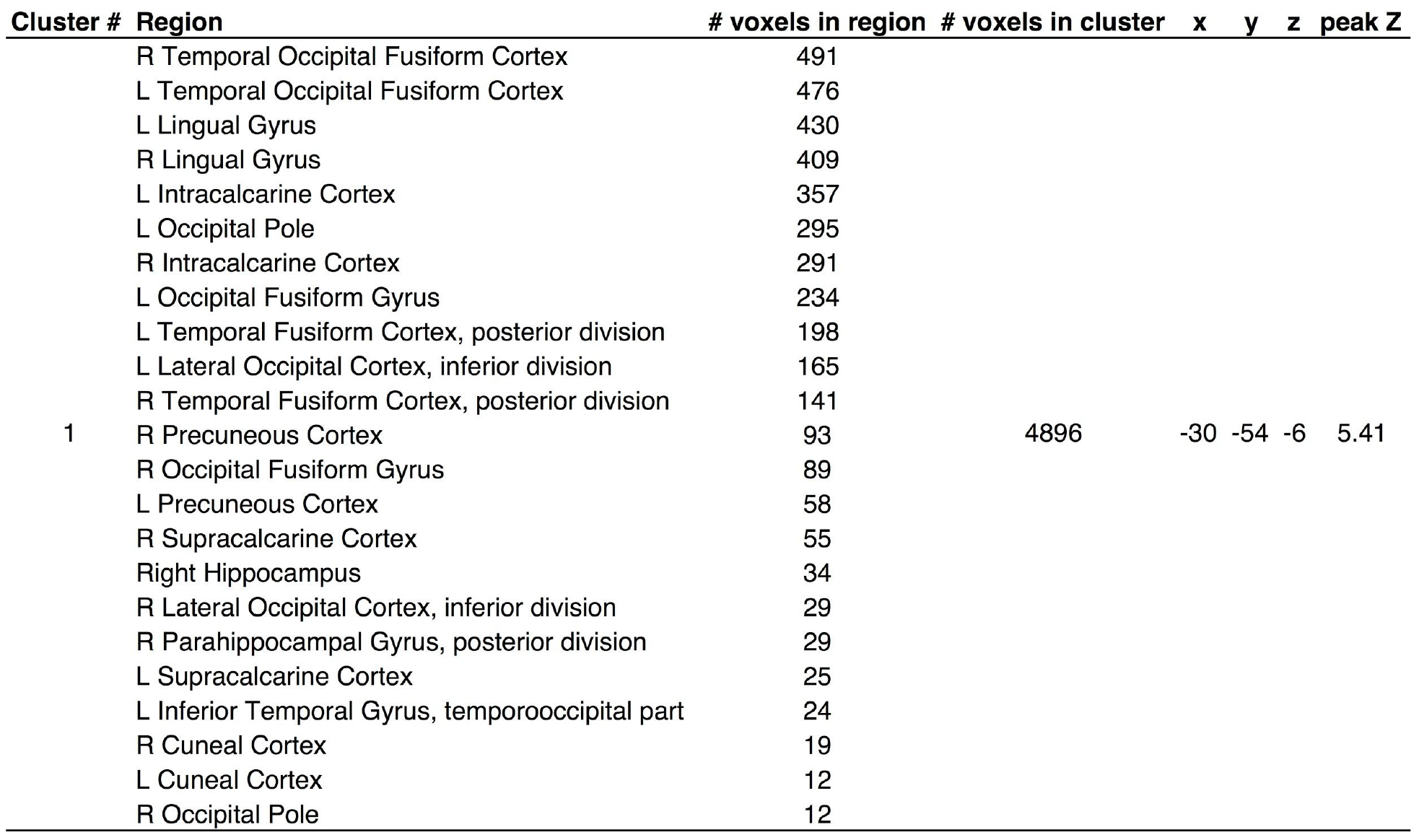
Activation table for map in Supplementary Figure 3b. (RT effect on BOLD for value-based greater than perceptual for trials with RTs that overlap in range between the two decision tasks)

**Supplementary Table 6.**
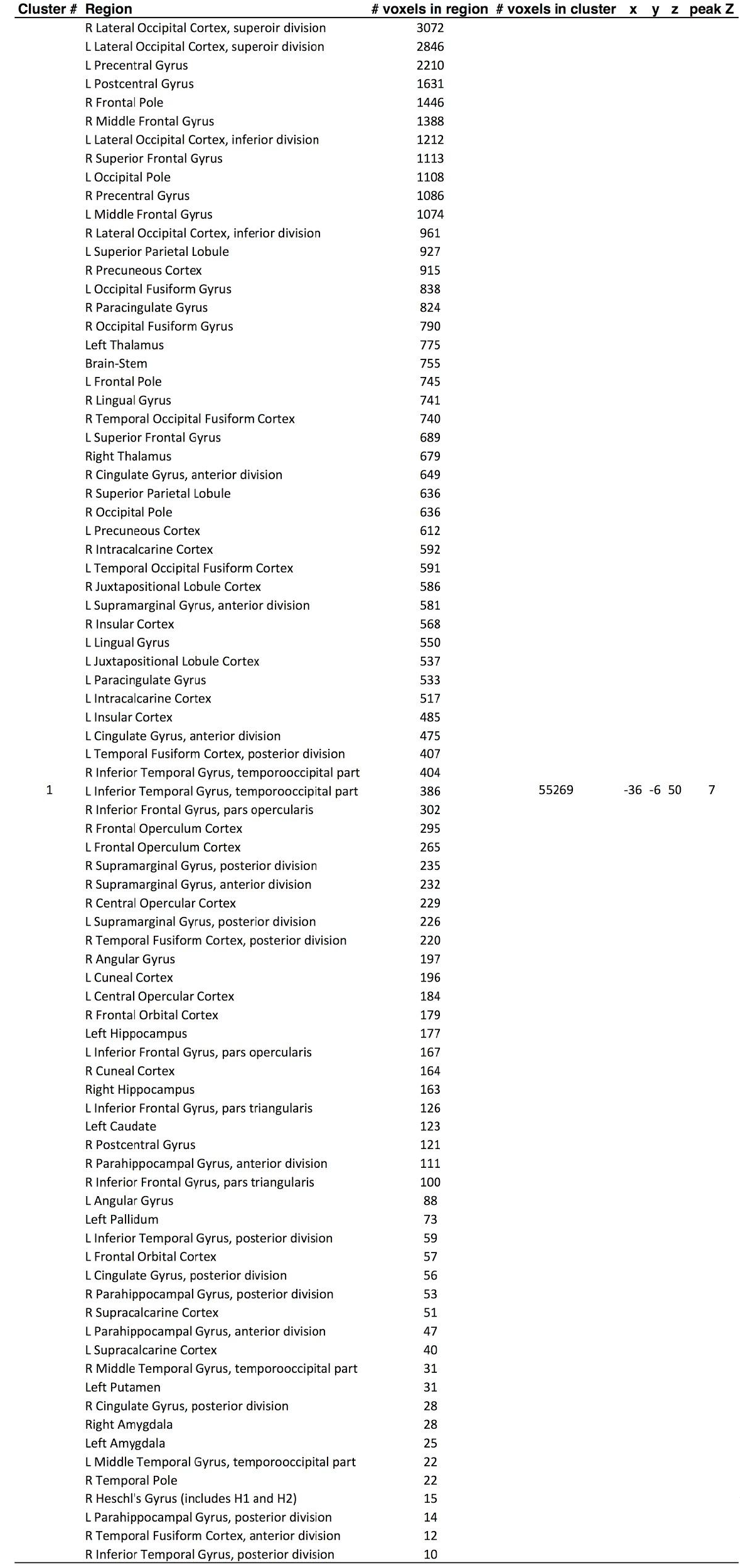
Activation table for map in Supplementary Figure 4a. (effect of value-based RT on BOLD)

**Supplementary Table 7.** Activation table for map in Supplementary Figure 4b. (effect of perceptual RT on BOLD)

| Cluster # | Region | # voxels in region | # voxels in cluster | x | y | z | peak Z |
| --- | --- | --- | --- | --- | --- | --- | --- |
| 1 | L Precentral Gyrus | 3111 | 54515 | -40 | -12 | 54 | 6.72 |
|  | R Lateral Occipital Cortex, superior division | 2731 |  |  |  |  |  |
|  | L Postcentral Gyrus | 2545 |  |  |  |  |  |
|  | L Lateral Occipital Cortex, superior division | 2259 |  |  |  |  |  |
|  | R Precentral Gyrus | 2108 |  |  |  |  |  |
|  | L Lateral Occipital Cortex, inferior division | 1553 |  |  |  |  |  |
|  | R Lateral Occipital Cortex, inferior division | 1496 |  |  |  |  |  |
|  | L Superior Parietal Lobule | 1396 |  |  |  |  |  |
|  | R Postcentral Gyrus | 1248 |  |  |  |  |  |
|  | R Superior Parietal Lobule | 1200 |  |  |  |  |  |
|  | R Occipital Fusiform Gyrus | 849 |  |  |  |  |  |
|  | L Occipital Fusiform Gyrus | 843 |  |  |  |  |  |
|  | L Occipital Pole | 818 |  |  |  |  |  |
|  | R Frontal Pole | 771 |  |  |  |  |  |
|  | R Occipital Pole | 746 |  |  |  |  |  |
|  | R Superior Frontal Gyrus | 741 |  |  |  |  |  |
|  | R Juxtapositional Lobule Cortex | 737 |  |  |  |  |  |
|  | R Cingulate Gyrus, anterior division | 727 |  |  |  |  |  |
|  | L Middle Frontal Gyrus | 723 |  |  |  |  |  |
|  | L Supramarginal Gyrus, anterior division | 683 |  |  |  |  |  |
|  | R Supramarginal Gyrus, anterior division | 680 |  |  |  |  |  |
|  | L Superior Frontal Gyrus | 597 |  |  |  |  |  |
|  | L Juxtapositional Lobule Cortex | 583 |  |  |  |  |  |
|  | R Middle Frontal Gyrus | 560 |  |  |  |  |  |
|  | L Frontal Pole | 552 |  |  |  |  |  |
|  | L Central Opercular Cortex | 545 |  |  |  |  |  |
|  | R Paracingulate Gyrus | 507 |  |  |  |  |  |
|  | Left Thalamus | 458 |  |  |  |  |  |
|  | R Temporal Occipital Fusiform Cortex | 449 |  |  |  |  |  |
|  | R Supramarginal Gyrus, posterior division | 444 |  |  |  |  |  |
|  | R Insular Cortex | 437 |  |  |  |  |  |
|  | Right Thalamus | 433 |  |  |  |  |  |
|  | L Cingulate Gyrus, anterior division | 430 |  |  |  |  |  |
|  | R Precuneous Cortex | 417 |  |  |  |  |  |
|  | L Insular Cortex | 410 |  |  |  |  |  |
|  | R Inferior Temporal Gyrus, temporooccipital part | 381 |  |  |  |  |  |
|  | L Paracingulate Gyrus | 380 |  |  |  |  |  |
|  | L Temporal Occipital Fusiform Cortex | 369 |  |  |  |  |  |
|  | R Central Opercular Cortex | 364 |  |  |  |  |  |
|  | L Parietal Operculum Cortex | 351 |  |  |  |  |  |
|  | R Middle Temporal Gyrus, temporooccipital part | 286 |  |  |  |  |  |
|  | L Supramarginal Gyrus, posterior division | 277 |  |  |  |  |  |
|  | L Precuneous Cortex | 253 |  |  |  |  |  |
|  | R Parietal Operculum Cortex | 249 |  |  |  |  |  |
|  | R Frontal Operculum Cortex | 246 |  |  |  |  |  |
|  | R Lingual Gyrus | 244 |  |  |  |  |  |
|  | L Frontal Operculum Cortex | 208 |  |  |  |  |  |
|  | L Inferior Temporal Gyrus, temporooccipital part | 195 |  |  |  |  |  |
|  | R Inferior Frontal Gyrus, pars opercularis | 193 |  |  |  |  |  |
|  | Brain-Stem | 172 |  |  |  |  |  |
|  | L Cingulate Gyrus, posterior division | 166 |  |  |  |  |  |
|  | L Planum Temporale | 165 |  |  |  |  |  |
|  | L Lingual Gyrus | 130 |  |  |  |  |  |
|  | R Cingulate Gyrus, posterior division | 118 |  |  |  |  |  |
|  | Left Putamen | 110 |  |  |  |  |  |
|  | L Heschl's Gyrus (includes H1 and H2) | 91 |  |  |  |  |  |
|  | R Planum Temporale | 85 |  |  |  |  |  |
|  | Right Hippocampus | 85 |  |  |  |  |  |
|  | R Angular Gyrus | 78 |  |  |  |  |  |
|  | L Intracalcarine Cortex | 77 |  |  |  |  |  |
|  | R Heschl's Gyrus (includes H1 and H2) | 75 |  |  |  |  |  |
|  | Right Pallidum | 73 |  |  |  |  |  |
|  | R Cuneal Cortex | 61 |  |  |  |  |  |
|  | L Middle Temporal Gyrus, temporooccipital part | 56 |  |  |  |  |  |
|  | Left Hippocampus | 55 |  |  |  |  |  |
|  | R Frontal Orbital Cortex | 54 |  |  |  |  |  |
|  | L Temporal Fusiform Cortex, posterior division | 48 |  |  |  |  |  |
|  | L Angular Gyrus | 47 |  |  |  |  |  |
|  | Left Pallidum | 46 |  |  |  |  |  |
|  | L Frontal Orbital Cortex | 42 |  |  |  |  |  |
|  | R Temporal Fusiform Cortex, posterior division | 41 |  |  |  |  |  |
|  | L Inferior Frontal Gyrus, pars opercularis | 39 |  |  |  |  |  |
|  | L Planum Polare | 36 |  |  |  |  |  |
|  | Right Putamen | 34 |  |  |  |  |  |
|  | L Inferior Temporal Gyrus, posterior division | 33 |  |  |  |  |  |
|  | R Inferior Frontal Gyrus, pars triangularis | 30 |  |  |  |  |  |
|  | R Inferior Temporal Gyrus, posterior division | 16 |  |  |  |  |  |
|  | L Inferior Frontal Gyrus, pars triangularis | 13 |  |  |  |  |  |
|  | R Temporal Pole | 13 |  |  |  |  |  |

**Supplementary Table 8.**
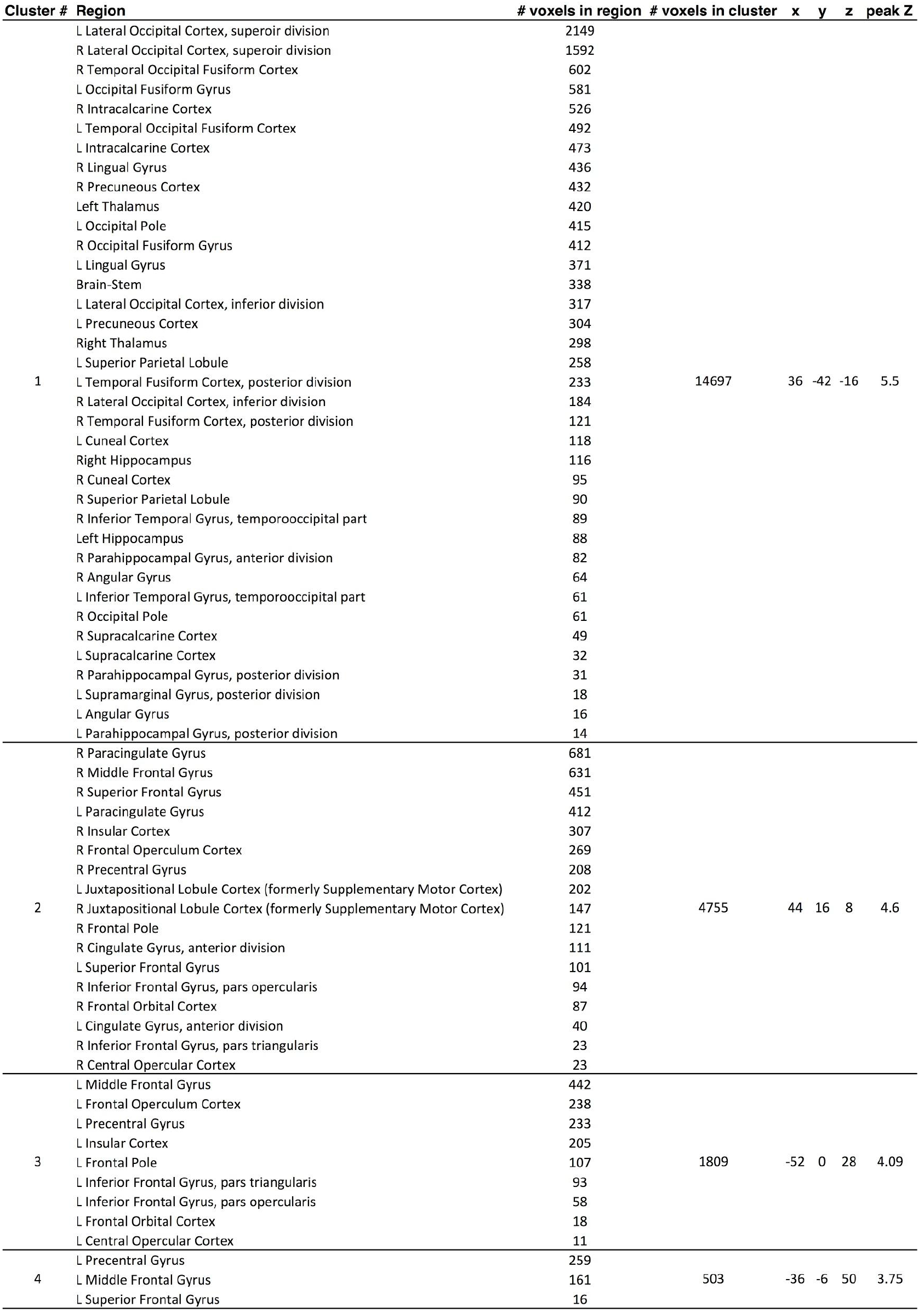
Activation table for map in Supplementary Figure 4c. (Value-based RT > Perceptual RT)

**Supplementary Table 9.** Activation table for map in Figure 3b. (PPI for slow > fast value-based decision trials with hippocampus seed)

| Cluster # | Region | # voxels in region | # voxels in cluster | x | y | z | peak Z |
| --- | --- | --- | --- | --- | --- | --- | --- |
| 1 | L Postcentral Gyrus | 411 | 1567 | -14 | -42 | 74 | 3.66 |
|  | R Postcentral Gyrus | 194 |  |  |  |  |  |
|  | R Precentral Gyrus | 184 |  |  |  |  |  |
|  | R Precuneous Cortex | 165 |  |  |  |  |  |
|  | L Superior Parietal Lobule | 127 |  |  |  |  |  |
|  | R Superior Parietal Lobule | 105 |  |  |  |  |  |
|  | L Precuneous Cortex | 56 |  |  |  |  |  |
|  | R Lateral Occipital Cortex, superior division | 47 |  |  |  |  |  |
|  | L Supramarginal Gyrus, anterior division | 31 |  |  |  |  |  |
|  | L Supramarginal Gyrus, posterior division | 30 |  |  |  |  |  |
|  | L Precentral Gyrus | 19 |  |  |  |  |  |
| 2 | R Frontal Orbital Cortex | 317 | 1328 | 34 | 28 | -20 | 3.89 |
|  | Right Putamen | 160 |  |  |  |  |  |
|  | R Insular Cortex | 158 |  |  |  |  |  |
|  | R Frontal Operculum Cortex | 145 |  |  |  |  |  |
|  | R Central Opercular Cortex | 69 |  |  |  |  |  |
|  | R Inferior Frontal Gyrus, pars triangularis | 56 |  |  |  |  |  |
|  | R Inferior Frontal Gyrus, pars opercularis | 39 |  |  |  |  |  |
|  | R Frontal Pole | 32 |  |  |  |  |  |
|  | Right Pallidum | 10 |  |  |  |  |  |
| 3 | R Superior Parietal Lobule | 300 | 696 | 34 | -46 | 50 | 3.42 |
|  | R Postcentral Gyrus | 213 |  |  |  |  |  |
|  | R Supramarginal Gyrus, anterior division | 106 |  |  |  |  |  |
|  | R Supramarginal Gyrus, posterior division | 11 |  |  |  |  |  |
| 4 | R Frontal Pole | 362 | 612 | 4 | 40 | 32 | 3.22 |
|  | R Paracingulate Gyrus | 135 |  |  |  |  |  |
|  | R Middle Frontal Gyrus | 25 |  |  |  |  |  |
|  | R Superior Frontal Gyrus | 13 |  |  |  |  |  |
| 5 | R Cingulate Gyrus, anterior division | 155 | 596 | 8 | -2 | 36 | 3.35 |
|  | R Juxtapositional Lobule Cortex | 89 |  |  |  |  |  |
|  | R Precentral Gyrus | 51 |  |  |  |  |  |
|  | L Juxtapositional Lobule Cortex | 49 |  |  |  |  |  |
|  | R Cingulate Gyrus, posterior division | 40 |  |  |  |  |  |
|  | Right Caudate | 19 |  |  |  |  |  |
|  | R Paracingulate Gyrus | 13 |  |  |  |  |  |
| 6 | R Temporal Occipital Fusiform Cortex | 231 | 581 | 34 | -58 | -18 | 3.34 |
|  | R Lateral Occipital Cortex, inferior division | 131 |  |  |  |  |  |
|  | R Inferior Temporal Gyrus, temporooccipital part | 66 |  |  |  |  |  |
|  | R Occipital Fusiform Gyrus | 21 |  |  |  |  |  |
|  | R Lingual Gyrus | 16 |  |  |  |  |  |
| 7 | L Lingual Gyrus | 109 | 578 | -10 | -62 | -12 | 3.61 |
|  | R Lingual Gyrus | 19 |  |  |  |  |  |
|  | L Occipital Fusiform Gyrus | 10 |  |  |  |  |  |
| 8 | L Insular Cortex | 211 | 550 | -30 | 20 | -8 | 3.27 |
|  | L Central Opercular Cortex | 48 |  |  |  |  |  |
|  | Left Putamen | 43 |  |  |  |  |  |
|  | L Frontal Operculum Cortex | 40 |  |  |  |  |  |
|  | L Frontal Orbital Cortex | 37 |  |  |  |  |  |
|  | L Precentral Gyrus | 29 |  |  |  |  |  |
|  | L Inferior Frontal Gyrus, pars opercularis | 19 |  |  |  |  |  |
| 9 | L Lateral Occipital Cortex, inferior division | 170 | 422 | -50 | -60 | -16 | 3.49 |
|  | L Inferior Temporal Gyrus, temporooccipital part | 140 |  |  |  |  |  |
|  | L Temporal Occipital Fusiform Cortex | 34 |  |  |  |  |  |
|  | L Inferior Temporal Gyrus, posterior division | 14 |  |  |  |  |  |
|  | L Temporal Fusiform Cortex, posterior division | 11 |  |  |  |  |  |

**Supplementary Table 10.** Activation table for map in Figure 4c. (effect of revalued value Vrev on BOLD)

| Cluster # | Region | # voxels in region | # voxels in cluster | x | y | z | peak Z |
| --- | --- | --- | --- | --- | --- | --- | --- |
| 1 | R Postcentral Gyrus | 357 |  |  |  |  |  |
|  | R Precentral Gyrus | 190 | 608 | 48 | -8 | 54 | 4.12 |
|  | R Superior Parietal Lobule | 47 |  |  |  |  |  |
| 2 | R Central Opercular Cortex | 142 |  |  |  |  |  |
|  | R Precentral Gyrus | 90 |  |  |  |  |  |
|  | R Superior Temporal Gyrus, posterior division | 56 |  |  |  |  |  |
|  | R Postcentral Gyrus | 28 | 513 | 42 | 0 | 18 | 3.2 |
|  | R Planum Temporale | 21 |  |  |  |  |  |
|  | R Heschl's Gyrus (includes H1 and H2) | 18 |  |  |  |  |  |
|  | R Superior Temporal Gyrus, anterior division | 12 |  |  |  |  |  |
| 3 | R Frontal Medial Cortex | 106 |  |  |  |  |  |
|  | L Frontal Medial Cortex | 96 |  |  |  |  |  |
|  | L Paracingulate Gyrus | 83 | 466 | 6 | 48 | -10 | 3.98 |
|  | R Paracingulate Gyrus | 61 |  |  |  |  |  |
|  | L Frontal Pole | 41 |  |  |  |  |  |
|  | R Frontal Pole | 32 |  |  |  |  |  |

